# Disordered regions of human eIF4B orchestrate a dynamic self-association landscape

**DOI:** 10.1101/2024.06.21.600094

**Authors:** Bikash Chandra Swain, Pascale Sarkis, Vanessa Ung, Sabrina Rousseau, Laurent Fernandez, Ani Meltonyan, V. Esperance Aho, Davide Mercadante, Cameron D. Mackereth, Mikayel Aznauryan

**Author notes:** Corresponding authors: Cameron D. Mackereth; Mikayel Aznauryan. these authors contributed equally (Bikash Chandra Swain, Pascale Sarkis, Vanessa Ung).

## Abstract

Eukaryotic translation initiation factor eIF4B is required for efficient cap-dependent translation, it is overexpressed in cancer cells, and may influence stress granule formation. Due to the high degree of intrinsic disorder, eIF4B is rarely observed in cryo-EM structures of translation complexes and only ever by its single structured RNA recognition motif domain, leaving the molecular details of its large intrinsically disordered region (IDR) unknown. By integrating experiments and simulations we demonstrate that eIF4B IDR orchestrates and fine-tunes an intricate transition from monomers to a condensed phase, in which large-size dynamic oligomers form before mesoscopic phase separation. Single-molecule spectroscopy combined with molecular simulations enabled us to characterize the conformational ensembles and underlying intra- and intermolecular dynamics across the oligomerization transition. The observed sensitivity to ionic strength and molecular crowding in the self-association landscape suggests potential regulation of eIF4B nanoscopic and mesoscopic behaviors such as driven by protein modifications, binding partners or changes to the cellular environment.

## Introduction

As their names suggest, eukaryotic translation initiation factors (or eIFs) play a critical role in the first stage of protein translation. At the heart of translation initiation in eukaryotes is the formation of two multi-factor assemblies: the cap-binding complex eIF4F formed in the vicinity of the mRNA 5’-end, and the 43S preinitiation complex (PIC) that forms in an mRNA-independent manner^1,2^. In terms of the main components, eIF4F consists of the cap-binding protein eIF4E, the RNA helicase eIF4A, and the large scaffolding protein eIF4G. These three factors ensure the recognition of 5’ mRNA 7-methylguanosine cap (via eIF4E) and the association to mRNA (via eIF4A and eIF4G)^3–5^. An additional role in unwinding 5’-proximal mRNA secondary structures relies on eIF4A, both as part of the eIF4F complex^3–5^, but also independently, as contributed by a second eIF4A possibly located at the mRNA entry channel of 40S small ribosomal subunit^6^. Moreover, the efficient mRNA unwinding requires the contribution of the initiation factor eIF4B (or the paralogue eIF4H) for full eIF4A activity and an increase in processivity^5,7–12^. The 43S PIC is a large assembly, consisting of the 40S ribosomal subunit, the ternary complex eIF2, GTP and Met-tRNAi, and several individual initiation factors (e.g. eIF1, eIF1A, eIF5) or multi-subunit complexes (such as the eIF3)^13^. Functionally, the 43S joins the eIF4F at the mRNA to form the 48S initiation complex^14^, which together with several auxiliary protein factors that again includes eIF4B (as well as eIF4H, eIF4A, DHX29, DDX3 and others) orchestrate the scanning of mRNA towards the start codon and start of protein synthesis^2,4,6,15^. A critical contribution to this mechanistic understanding of the eukaryotic translation initiation resulted from a burst of high-resolution structural characterizations of various translation initiation complexes and sub-complexes^6,14,16–19^. However, due to the nature of cryo-EM and crystallography approaches, these experimental advances nevertheless omit the characterization of the constituent proteins and protein regions that are intrinsically disordered.

Specifically within this latter category of proteins is eIF4B, the major portion of which is predicted to be intrinsically disordered (Fig. 1a,b). In addition to the specific activation of eIF4A detailed above, eIF4B is generally required for efficient control of cell survival and proliferation^20–22^ as it selectively regulates a set of proliferative and prosurvival mRNAs that encompass long structured 5′ untranslated regions (UTRs)^20,23^. In this context, eIF4B was shown to be overexpressed in certain cancer cells and the reduction of eIF4B expression is sufficient to decrease synthesis of proteins associated with enhanced tumor cell survival^24^. Thus, eIF4B is considered as a potential anti-cancer target^22^. eIF4B has also been reported to localize into stress granules^25,26^, and the reduction of cellular eIF4B levels was shown to affect stress granule formation^27^.

Human eIF4B consists of 611 residues and is composed of a folded RNA recognition motif (RRM) domain preceded by a short unstructured tail^28^, and an approximately 400 residue intrinsically disordered region (IDR) downstream the RRM (Fig. 1a,b). As the single folded domain present in eIF4B, the RRM domain has been previously studied at the atomic level^28^. Recently, we have also characterized the atomic details of the C-terminal region (CTR) of eIF4B, encompassing residues 333 to 611, and experimentally confirmed that this region is indeed highly intrinsically disordered^29^. Nonetheless, we could also identify a short helical segment that coincides with the previously mapped arginine-rich motif (ARM) implicated in RNA binding^11,30^ (Fig. 1b).

Another notable segment of the IDR encompasses a defined low-complexity region enriched in aspartate, arginine, tyrosine and glycine residues (the DRYG-rich region, Fig. 1b,c)^31,32^. Initially this region consisted of residues 164-356,^31^ but later it was refined and mapped to residues 214-327^11,32^, to precisely bracket the cluster of tyrosine and glycine residues (Fig. 1c). In some deletion variants, the DRYG-rich region was alternatively defined to extend further towards the C-terminus, covering the residues 213-340.^30^ This DRYG-rich region was shown to facilitate the direct interaction between eIF4B and eIF3a, which is believed to mediate the association of the mRNA with 40S^32^. At the same time, the deletion of the DRYG-rich region from the full-length eIF4B reduces the RNA binding affinity^30^, either because this region has intrinsic RNA affinity or it stabilizes the RNA interaction with the RNA-binding region ARM^30^. The key feature of the DRYG-rich region is its ability to promote the self-association of eIF4B, both *in vitro* and *in vivo*^32^. Moreover, the N-terminal ∼210 residue portion of eIF4B that encompasses the RRM domain and its N-terminal tail is dispensable for self-association^32^. Notably, the removal of the DRYG-rich region leads to a loss of global translation inhibition associated with the overexpression of eIF4B in certain mammalian cells^30,33^, suggesting that DRYG-driven self-association of eIF4B might be crucial for its function in translation regulation. The exact links between these observations however remain unexplained, partly due to the lack of description of molecular principles underlying the eIF4B self-association.

Thus, with a goal of understanding the mechanistic bases of eIF4B self-association we focused on the C-terminal IDR of eIF4B, starting from the DRYG-rich region up to the complete C-terminus and performed a detailed characterization of structural dynamics and interactions using a combination of single-molecule FRET spectroscopy (smFRET) and nuclear magnetic resonance (NMR) spectroscopy experiments, coarse-grained Langevin dynamics simulations and phase separation assays. By integrating the atomic and molecular information at different scales, we obtained a comprehensive picture of factors defining the molecular behavior of eIF4B IDR across several regimes of the self-association landscape, ranging from monomers to oligomers and condensates.

**Fig. 1:**
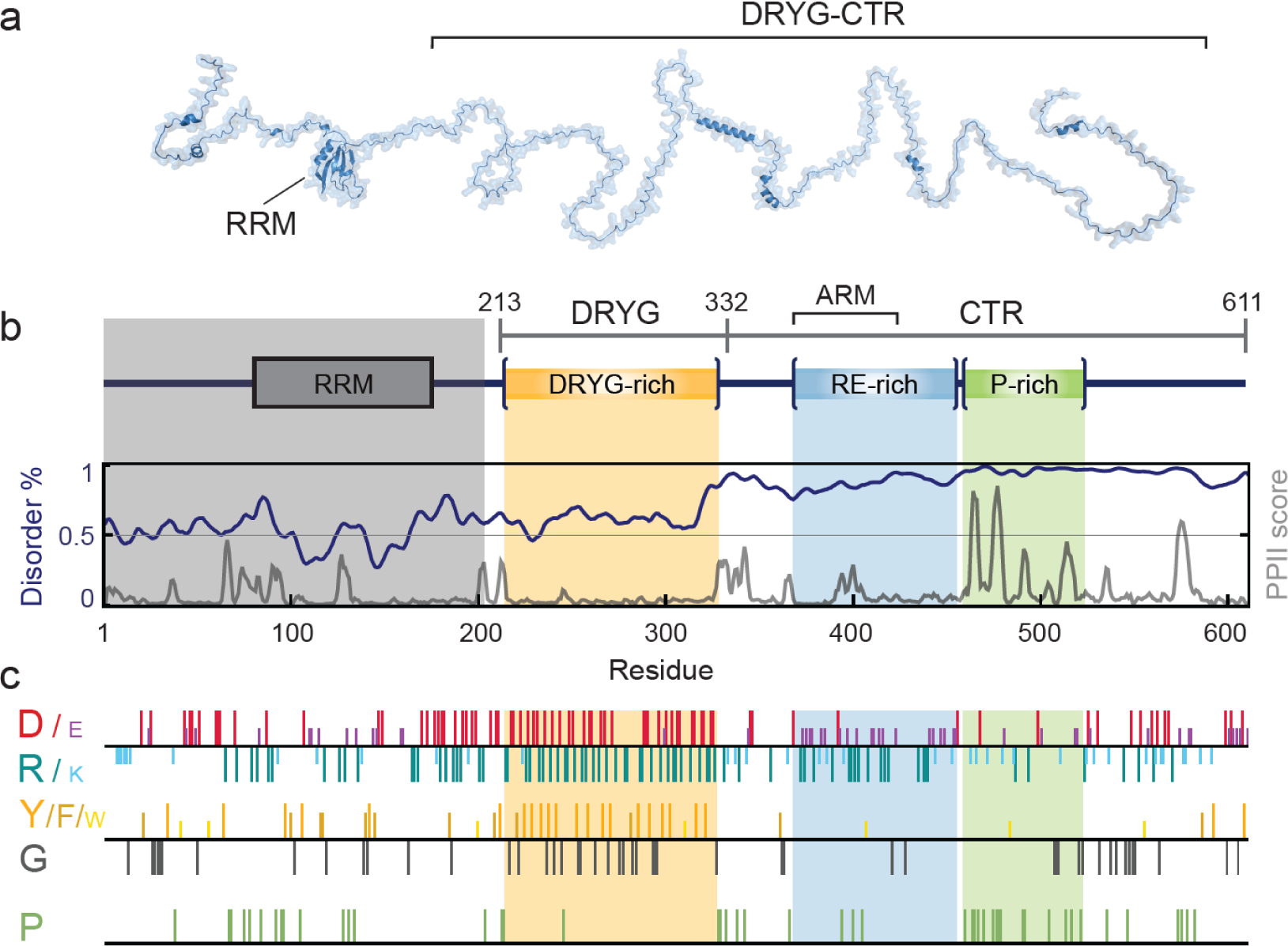
An overview of sequence and structural characteristics of eIF4B. **a** Cartoon representation of human eIF4B based on the RRM domain structure (PDB: 2J76) and predicted IDR structural propensity from AlphaFold2. **b** Schematic domain representation of human eIF4B, together with a disorder prediction plot based on IUPred3^34^ (dark blue) and the prediction of polyproline II secondary structure propensity (gray)^35^. The structured RRM domain and short disordered N-terminal tail are shadowed in gray. The boundaries of protein constructs used in this work are also indicated. **c** Amino acid distribution plots for negative (D and E), positive (R and K), aromatic (Y, F and W), glycine (G) and proline (P) residues, particularly highlighting clustering of these residues within several subregions, i.e. DRYG-rich region (residues 214-327), RE-rich region (residues 367-455) and the P-rich region (residues 460-522).

## Results

### DRYG-CTR undergoes homotypic phase separation

As the DRYG-rich segment in the IDR has a reported role in eIF4B self-association, we first focused on this fragment of eIF4B for detailed investigation. The minimal DRYG protein fragment (residues 214-332) was purified from inclusion bodies, where it exclusively localized during recombinant expression. In line with this observation, the DRYG-rich fragment was only soluble in the presence of guanidine hydrochloride (GdmCl) and only within a limited protein concentration range (up to ∼10 μM in 0.5 M GdmCl). At higher concentrations the purified construct again underwent phase separation, effectively prohibiting characterization of the minimal DRYG-rich region under native conditions. From literature, the start of the DRYG-rich region is well defined (i.e. residue 214)^11,32^ whereas the end residue is more vaguely specified and had been suggested to span beyond residue 327.^30,31^ The easiest option therefore was to extend the protein construct from the DRYG-rich region towards the final C-terminal end of the protein (residues 213-611), thus encompassing the almost complete C-terminal IDR of eIF4B in a construct we term DRYG-CTR (Fig. 1a,b).

Already at the stage of protein purification, we noticed that DRYG-CTR also undergoes phase separation, albeit to much lesser degree as compared to the minimal DRYG-rich fragment and thus was readily amenable to further characterization. Differential interference contrast (DIC) and fluorescence imaging confirmed the formation of microscopic phase-separated droplets, which undergo the fast fusion characteristic of liquid-like behavior (Fig. 2a,b). To map the condensation properties of DRYG-CTR, we first quantified the protein concentration in dilute (*c*_sat_) and dense (*c*_dense_) phases at different temperatures. This concentration window defines the DRYG-CTR phase separation regime, outside of which the protein is easily soluble and miscible in solution (Fig. 2c). The resulting temperature-dependent phase diagram reveals an increase of phase separation propensity at lower temperatures, following a characteristic upper critical solution temperature (UCST) behavior. At the same time, the phase separation of DRYG-CTR is favored in low ionic strength buffers, e.g. 20 mM sodium phosphate (NaP), 50 mM NaCl, pH 7.0 (Fig. 2c), and is largely suppressed (*c*_sat_ > 1.5 mM) at NaCl concentrations above 150 mM.

### eIF4B IDR is largely devoid of stable secondary structure

To proceed with structural characterization of DRYG-CTR, we next assessed the degree of protein secondary structure by circular dichroism (CD), under conditions suppressing protein phase separation (Fig. 2d). The CD spectrum shows a large minimum at 202 nm and small ellipticity signal at 220 nm, which is consistent with low secondary structure content as expected for disordered proteins. The small contribution of ellipticity at 220 nm is likely originating from transient short helical motifs at the CTR suggested by our previous NMR data^29^ (Supplementary Fig. 1).

The ^1^H–^15^N heteronuclear single quantum coherence (HSQC) spectrum of ^15^N-labeled DRYG-CTR is useful to probe at the residue level the nature of protein behavior. Under a variety of conditions, the only backbone amides that are observed appear as crosspeaks in the narrow ^1^H chemical shift range from 8.0-8.5 ppm, which is characteristic of disordered peptides (Fig. 2e). Nevertheless, at low ionic strength (50 mM NaCl), the ^15^N-HSQC spectrum of DRYG-CTR displays only a few clear resonance peaks, which account for only a small portion of residues from the full construct. In particular, the glycine region in the spectrum is easy to analyze (circled in Fig. 2e), and only 10 out of the total 36 expected glycine backbone amide crosspeaks are observed. Line-broadening (that is, loss of crosspeak intensity) can be caused by several factors including dynamics, heterogeneous folded states or formation of large oligomers, and in this case, it is most likely consistent with the condensation of the protein that is observed in low ionic strength. With an increase in ionic strength (300 mM NaCl) this number approximately doubles the observed glycine amide crosspeaks (Fig. 2e, middle spectrum.). The appearance of more peaks follows the reduced ability to phase separate observed at higher salt concentrations, which is expected to result in smaller protein assemblies. As a control experiment, the construct was fully destabilized upon addition of 1 M guanidinium hydrochloride (GdmCl) which should prevent essentially all inter- and intramolecular interactions. Indeed, this ^15^N-HSQC spectrum has equal intensity sharp amide peaks for what appears to be the entire protein sequence, as specifically evidenced by observation of all 36 glycine backbone amides (Fig. 2e, right spectrum).

To return to the DRYG-CTR construct at moderate ionic strength, the number and position of observed amide crosspeaks is remarkably similar to the pattern observed for the isolated CTR^29^. An overlay of the DRYG-CTR and CTR spectra at 150 mM NaCl highlights this similarity (Fig. 2f) and suggests that the visible residue crosspeaks all derive from the CTR region, and that the DRYG residue amides are in contrast all line-broadened. Our previous chemical shift assignment of human eIF4B CTR allows for a higher-resolution analysis, and it appears that not only the DRYG region, but most of the N-terminal third of the CTR displays markedly different behavior in the DRYG-CTR sample (residues 332-450, such as seen for G362, G363) (Fig. 2g). There also appears to be a small segment in the C-terminus from residue 580–600 (such as A590, A594) that also has more peak-broadening in the DRYG-CTR sample. Taken together, it appears that the DRYG-CTR construct spectra are consistent with the formation of a large, possibly heterogenous, assembly centered on the DRYG region but also encompassing to a lesser degree the neighboring region of CTR and a small C-terminal segment. Unfortunately, the NMR data also demonstrate that increased insight in the molecular nature of DRYG-CTR will require complementary biophysical techniques.

**Fig. 2:**
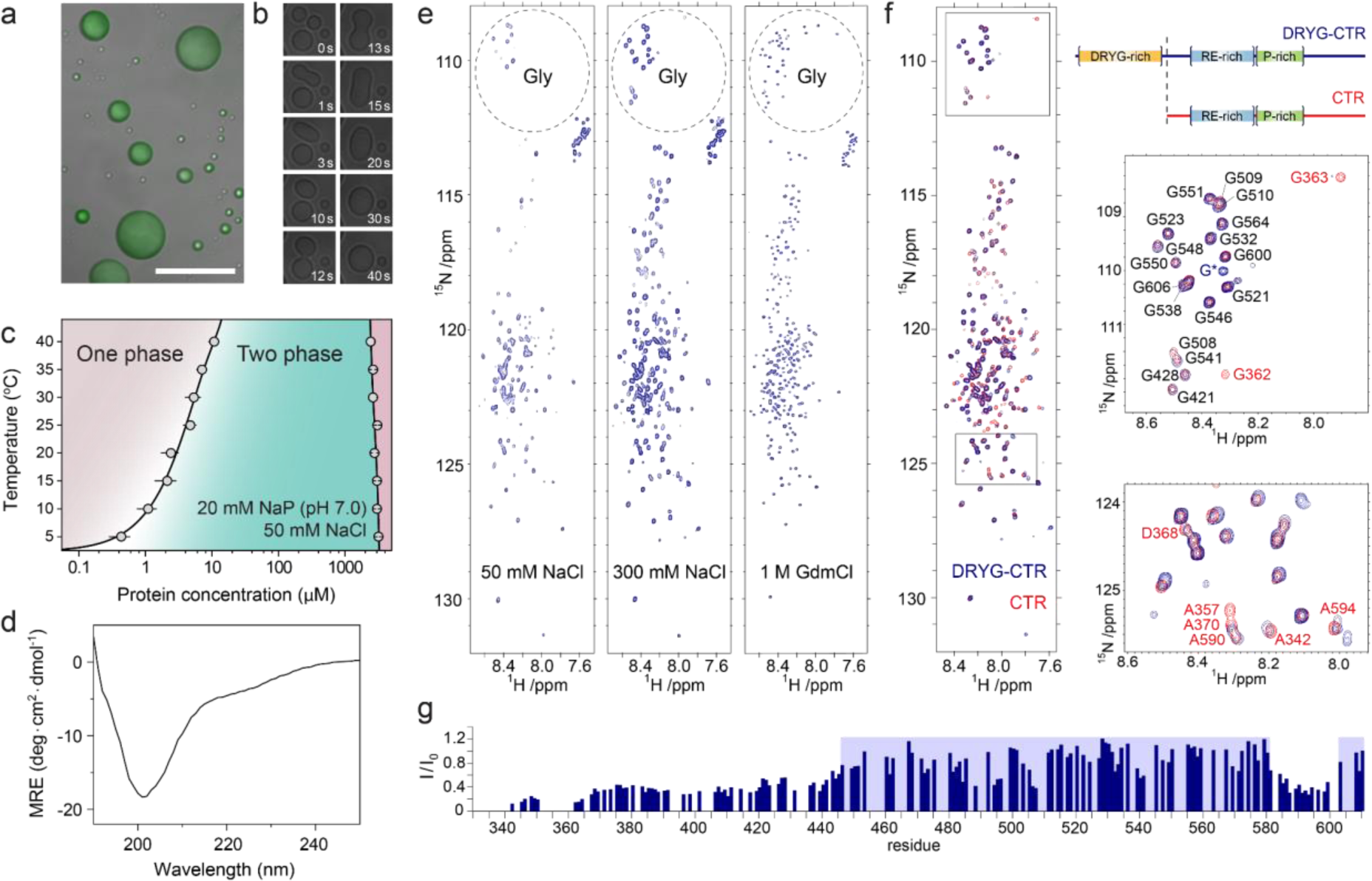
The structural and phase separation behavior of eIF4B IDR. **a** An overlay of DIC and fluorescence images of liquid-like droplets forming upon phase separation of DRYG-CTR. The scale bar indicates 20 μm. **b** A time course of DIC images showing rapid fusion of protein droplets. **c** Phase diagram of DRYG-CTR in 20 mM NaP, 50 mM NaCl (pH 7.0), showing the temperature dependence of *c*_sat_ (left branch) and *c*_dense_ (right branch). **d** Far-UV CD spectrum of DRYG-CTR in 20 mM NaP, 75 mM NaCl (pH 7.0), highlighting the low secondary structure content in DRYG-CTR. **e** ^1^H,^15^N-HSQC spectra of DRYG-CTR in 20 mM NaP (pH 7.0) buffer with 50, 300 mM NaCl and 1 M GdmCl. The dashed circles highlight resonance peaks for glycine residues. **f** The overlay of HSQC spectra for DRYG-CTR (blue) and CTR (red) constructs in 20 mM NaP, 150mM NaCl (pH 7.0). The right panel shows the schematics of DRYG-CTR and CTR constructs, together with closeup segments of the overlaid HSQC spectra. **g** The ratio of peak intensities for DRYG-CTR and CTR constructs, the blue shaded areas indicate the protein regions with comparable peak heights between the two constructs.

### Sequence bias defines conformational subregions of distinct compactness and dynamics

Although the NMR analysis confirms the disordered nature of the CTR, as well as general segments involved in self-association, the high-resolution data was restricted to residues outside of the DRYG region. To overcome this limitation in the NMR analysis we opted to characterize the conformational behavior and structural dynamics of DRYG-CTR using smFRET, as this technique is amenable to observation of defined subregions in a wide range of protein states, even in conformationally heterogenous samples. For this purpose, we created four different fluorescently labeled DRYG-CTR constructs: DRYG-CTR_P213C-P332C_, DRYG-CTR_P332C-C457_, DRYG-CTR_C457-G523C_ and DRYG-CTR_G523C-Y609C_ (Fig. 3a). We selected the labeling positions to ensure that each labeled construct probes distinct segments of specific sequence biases in the context of complete DRYG-CTR. In this context, the residues 213 to 332 bracket the low-complexity DRYG-rich region. Additionally, we defined the segment of residues 367 to 455, which partially overlaps with previously identified ARM region, and termed it the RE-rich region, due to its enrichment in alternating arginine and aspartate residues. Downstream to this region we defined a segment of residues 460 to 522, which we termed P-rich region, as it contains a large cluster of prolines. Additionally, we note the characteristic large negative net charge of the far C-terminal region (residues 523-611), particularly due to the cluster of glutamate and aspartate residues at the end of the protein. Based on these constructs, we then performed a large set of smFRET and nanosecond fluorescence correlation spectroscopy (nsFCS) experiments to characterize the corresponding protein regions of DRYG-CTR.

Starting with smFRET analysis of the DRYG subregion, the transfer efficiency (E) histograms of DRYG-CTR_P213C-P332C_ (Fig. 3b and Supplementary Fig. 2) show single relatively narrow peaks consistent with a fast-interconverting ensemble of disordered conformations. At the same time, the analysis of fluorescence lifetimes reveals that the normalized donor lifetime versus transfer efficiency histograms systematically deviate from the static diagonal line, and instead are centered along the curved line. This behavior is expected for a broad distance distribution of disordered proteins. Considering the large sequence separation between the two fluorophores (120 amino acids), the transfer efficiency histogram of DRYG-CTR_P213C-P332C_ is centered at a surprisingly high mean transfer efficiency, which indicates that this labeled segment displays an ensemble of very compacted disordered conformations. The conformation of the probed region is only slightly affected by ionic strength, as evidenced by a small increase of mean distance (*R*_DA_) between the fluorophores upon increasing concentration of NaCl. In contrast, the transfer efficiency peak monotonically shifts toward lower transfer efficiencies with increasing GdmCl concentration (Fig. 3b,c and Supplementary Fig. 3), indicating a continuous expansion of this segment, similar to other unfolded proteins and IDPs^36–38^.

The nsFCS measurements with DRYG-CTR_P213C-P332C_ enabled us to quantify the long-range distance dynamics between the two fluorophores bracketing this protein segment (Fig. 3d and Supplementary Fig. 4). In the absence of GdmCl, we observed chain reconfiguration times (*τ_r_*) in the range of 150 ns that is within a typical timescale for IDPs^39^, albeit reflecting overall reduced fluctuations of the polypeptide chain. The relatively small *R*_DA_ and longer *τ_r_* values for DRYG-CTR_P213C-P332C_ indicate a large contribution of transient intramolecular contacts within this segment that compact the conformational ensemble and slow down the chain dynamics of the corresponding DRYG-rich region. As both the *R*_DA_ and the *τ_r_* values are practically unaltered by the increase of the ionic strength we conclude that these intramolecular interactions are not electrostatic, and are likely to be driven by cation-π (between the arginine and tyrosine residues) and π-π stacking interactions (between the tyrosine residues). Increasing the GdmCl concentration results in decrease of the *τ_r_* values, consistent with GdmCl-induced disruption of transient intramolecular interactions within the DRYG-rich region and thus faster disordered chain fluctuations upon its expansion (Fig. 3c,d).

Altogether, our smFRET data confirms that the DRYG-rich region adopts relatively compact but nevertheless disordered and flexible conformations, with overall reduced chain dynamics.

Similar to DRYG-CTR_P213C-P332C_, the transfer efficiency histograms of the DRYG-CTR_P332C-C457,_ DRYG-CTR_C457-G523C_ and DRYG-CTR_G523C-Y609C_ also show single narrow peaks centered at intermediate E values, meanwhile the fluorescence lifetimes cluster above the diagonal line, which collectively confirm the more expanded and flexible nature of these labeled regions (Fig. 3e and Supplementary Fig. 2,3). The chain reconfiguration times, *τ_r_*, from nsFCS measurements are in the range of ∼100 ns (Fig. 3d and Supplementary Fig. 4), which is typical for disordered proteins of comparable lengths^38,39^. In contrast to the DRYG-rich region, these labeled segments are sensitive to the ionic strength, expanding to various degrees upon the increase of NaCl concentration (Fig. 3f). This behavior indicates the contribution of intramolecular electrostatic contacts, due to a large fraction of oppositely charged residues (Fig. 1c), which weaken upon electrostatic screening at higher ionic strengths. A similar expansion is also observed for these segments upon increasing GdmCl concentrations (Fig. 3f and Supplementary Fig. 3). It is remarkable that the dimensions of probed regions do not scale with their amino acid sequence length, as it would be expected for an ideal disordered chain in the absence of any intra-chain interactions (Fig. 3g). As already mentioned, the main outlier is the DRYG-CTR_P213C-P332C_, which has a scaling exponent (ν =0.43) characteristic of compact coils. For the other three constructs we observe larger scaling exponents (ν >0.5) consistent with more expanded conformations of corresponding probed regions. It is worth mentioning that DRYG-CTR_C457-G523C_, which possesses larger scaling exponent (ν =0.58) compared to DRYG-CTR_P332C-C457_ and DRYG-CTR_G523C-Y609C_, which can be attributed to the presence of a large cluster of proline residues (Fig. 1b). The proline-rich region is expected to increase the stiffness of the polypeptide chain and promote transient polyproline-II helix formation (Fig. 1b), thus resulting in more expanded conformation of this region^40,41^. The stiffness of the DRYG-CTR_C457-G523C_ is also evident from the relatively larger values of *τ_r_*, (e.g. compared to DRYG-CTR_G523C-Y609C_), as it would also be expected to scale with the chain length in case of similar degree of internal friction^37,42^. Scaling exponents of these three regions systematically increase upon addition of GdmCl and cluster around the ν =0.59±0.01, as expected for largely expanded disordered chains in the presence of denaturants. Here as well, the DRYG-CTR_P213C-P332C_ possesses a smaller scaling exponent, indicating that the corresponding protein region is yet more compact compared to the ideal chain, likely due to a fraction of intrachain arginine and tyrosine mediated cation-π and π-π contacts that persist even at high GdmCl concentrations.

Despite being largely unstructured, the eIF4B IDR therefore does not exhibit a uniform conformational behavior and plasticity. Different regions instead possess a variable degree of compactness and dynamics, primarily dictated by the amino acid composition of these segments and underlying intrachain contacts. These two features also define the protein response to changes of ionic strength, with the DRYG-rich region essentially unaffected by an increase of salt concentration, whereas the CTR expands upon screening of intrachain charge-charge interactions.

**Fig. 3:**
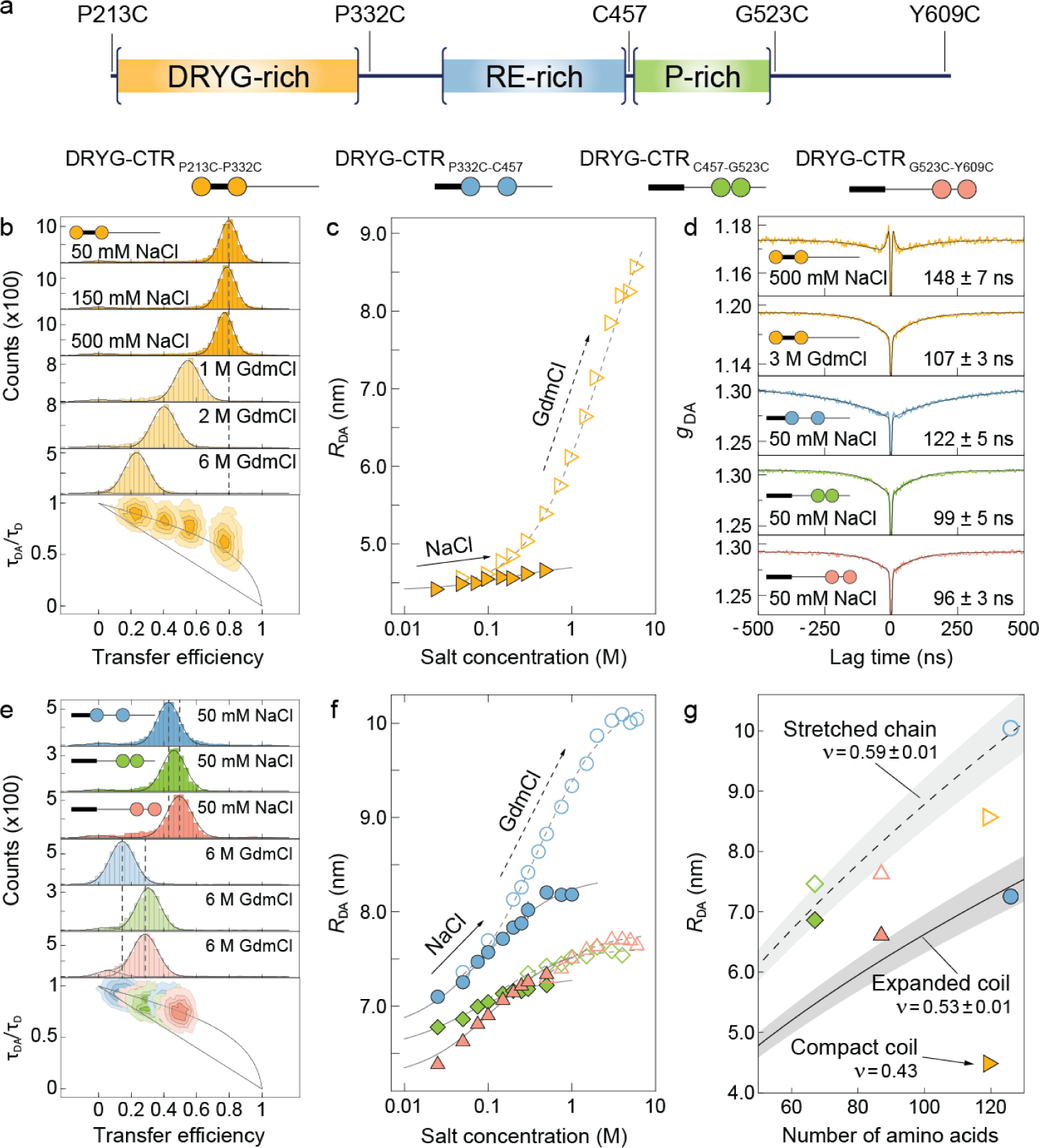
Different regions of eIF4B IDR have distinct compactness and dynamics. **a** A schematic representation of the DRYG-CTR construct, highlighting the pairs of cysteine residues used for fluorescent labeling of the protein. The bottom panel shows icons for four fluorescently labeled protein constructs used in smFRET experiments. **b** Transfer efficiency histograms of DRYG-CTR_P213C-P332C_ with different concentrations of NaCl and GdmCl. The dashed lines indicate mean transfer efficiency at 50 mM NaCl. The bottom panel shows the histograms of normalized donor fluorescence lifetime against transfer efficiency, acquired under the same conditions as the transfer efficiency histograms. The diagonal and curved lines indicate the donor lifetime - transfer efficiency dependence expected for fixed distance and broad distance distribution, respectively. **c** The mean inter-residue distance, *R*_DA_, for DRYG-CTR_P213C-P332C_ as a function of NaCl (filled symbols) and GdmCl (open symbols) concentration. The solid and dashed lines show the fits to an empirical binding isotherm. **d** nsFCS donor-acceptor correlation curves for DRYG-CTR_P213C-P332C_ acquired at 500 mM NaCl and 3 M GdmCl, and for DRYG-CTR_P332C-C457,_ DRYG-CTR_C457-G523C_ and DRYG-CTR_G523C-Y609C_ at 50 mM NaCl. The solid lines are fits used to determine the *τ_r_* reconfiguration times, as indicated (see Methods for details). **e** Transfer efficiency histograms of DRYG-CTR_P332C-C457,_ DRYG-CTR_C457-G523C_ and DRYG-CTR_G523C-Y609C_ at 50 mM NaCl and 6 M GdmCl. The dashed lines indicate mean transfer efficiency for DRYG-CTR_P332C-C457_ and DRYG-CTR_G523C-Y609C_ in the presence of NaCl and GdmCl. The bottom panel shows histograms of normalized donor fluorescence lifetime against transfer efficiency, acquired under the same conditions as the transfer efficiency histograms. **f** The mean inter-residue distance, *R*_DA_, for DRYG-CTR_P332C-C457,_ DRYG-CTR_C457-G523C_ and DRYG-CTR_G523C-Y609C_ as a function of NaCl (filled symbols) and GdmCl (open symbols) concentration. The solid and dashed lines show the fits to an empirical binding isotherm. **g** The dependence of mean inter-residue distance for DRYG-CTR_P213C-P332C_, DRYG-CTR_P332C-C457,_ DRYG-CTR_C457-G523C_ and DRYG-CTR_G523C-Y609C_ as a function of sequence separation, at 50 mM NaCl (filled symbols) and 6 M GdmCl (open symbols). The solid and dashed lines (and corresponding gray bands) indicate the expected *R*_DA_ - *N*_aa_ dependence with scaling exponent ν=0.53±0.01 and ν =0.59±0.01, respectively.

### eIF4B IDR self-associates into large-size oligomers driven by DRYG-rich region

As eIF4B was previously reported to form simple homodimers mediated by DRYG-rich region^32^, we wondered if DRYG-CTR may be able to self-associate at sub-saturation concentrations independently of phase separation (Fig. 2a-c), and in particular at conditions in which phase separation is not observed. We therefore used single-molecule fluorescence correlation spectroscopy (FCS) to probe the sub-nanomolar concentrations of fluorescently labeled monomeric DRYG-CTR upon titration with unlabeled protein (Fig. 4a). Upon increasing unlabeled protein concentrations, we observed a shift of FCS curves towards longer times, consistent with DRYG-CTR self-association and molecular size increase for diffusing protein species (Fig. 4b). Similar experiments with the CTR construct show no change in the protein diffusion rate, confirming the essential role of DRYG-rich region in oligomerization (Supplementary Fig. 5). Fitting FCS data allowed us to extract the average diffusion rates of diffusing protein species. The relative diffusion rates gradually decrease upon self-association at increasing protein concentrations (Fig. 4c). Surprisingly, the observed drop of diffusion rates extends beyond what would be anticipated for simple dimers, reaching a plateau at values corresponding to oligomers with an apparent average size of >10-mers. Thus, the eIF4B IDR does not undergo simple dimerization, as previously proposed^32^, but forms rather large oligomers.

Interestingly, the observed average size of the oligomers is slightly dependent on ionic strength, plateauing at average oligomer size of ∼15-mers in the presence of 75-150 mM NaCl, but dropping to ∼10-mers at higher ionic strength. The midpoint of monomer to oligomer transition curves shown in Fig. 4c reflects the apparent dissociation constant, *K*_D_, of oligomers, which decreases upon lowering the NaCl concentration (Fig. 4d), indicating an increased DRYG-CTR association affinity. Interestingly, the change of dissociation constants with ionic strength is very modest and the analysis of the NaCl dependence of *K*_D_s, reveals a relatively small number of counterions (Δ*n* = 1.86 ± 0.04) released upon protein self-association (Fig. 4d). This points towards a weak electrostatic nature of DRYG-DRYG mediated protein association and implies that DRYG-CTR intermolecular interactions are presumably dictated by cation-π and π-π contacts, similar to the intramolecular interactions observed within DRYG-rich region.

**Fig. 4:**
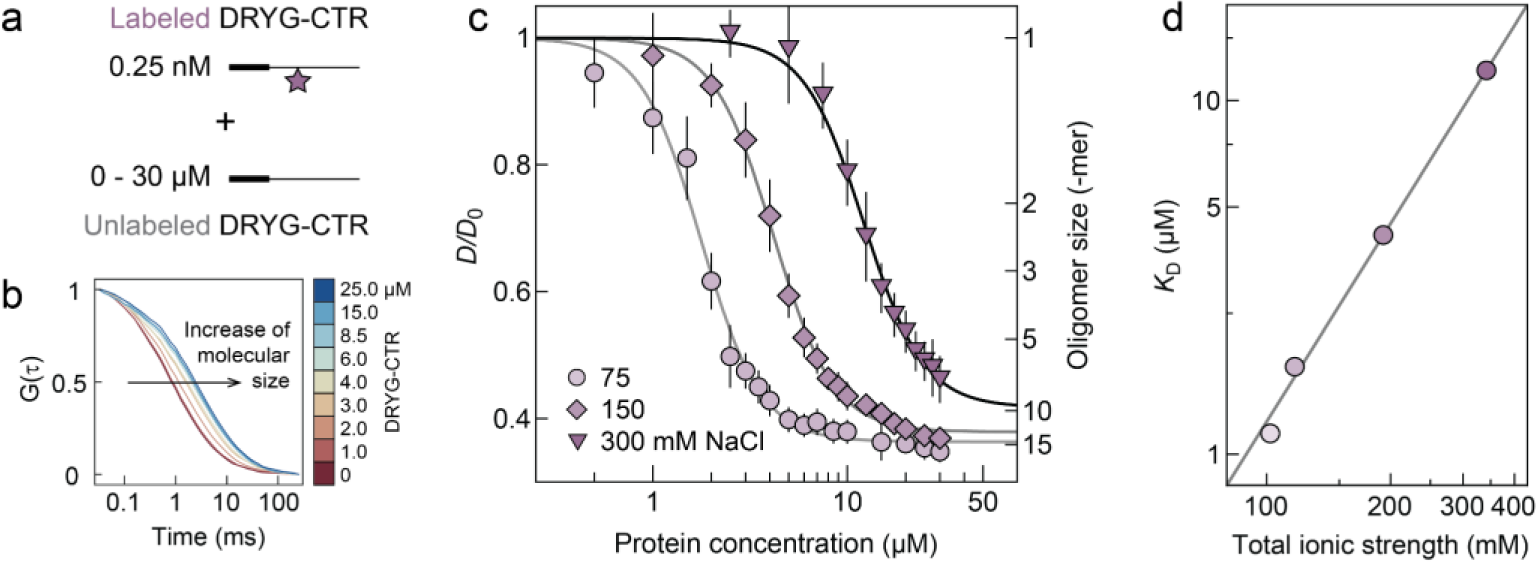
eIF4B IDR undergoes self-association and forms large-size oligomers, with weak dependence on ionic strength. **a** A schematic of FCS oligomerization assay, based on a titration of sub-nanomolar labeled DRYG-CTR with micromolar concentrations of unlabeled protein. **b** A set of FCS curves at 75 mM NaCl obtained in the presence of increasing protein concentration show a clear shift towards higher times, indicating a growing size of diffusing species. **c** The dependence of relative diffusion rates (normalized to the diffusion rate of monomeric protein) on protein concentration, acquired in buffers containing 75, 150 and 300 mM NaCl, indicates a reduction of protein association affinity with increasing buffer ionic strength. The solid lines show global fits to the Hill equation. **d** The dependence of apparent dissociation constant on buffer ionic strength. The black solid line is a fit with the Lohman– Record model^43^.

### DRYG-CTR oligomers are disordered and dynamic complexes

Although FCS experiments allowed us to extract substantial details on DRYG-CTR self-association behavior, they did not provide information on the structural and dynamic aspects underlying the oligomerization process. To address this aspect, we took advantage of the double-labeled DRYG-CTR constructs, and performed smFRET oligomerization assays probing transfer efficiency histograms of labeled DRYG-CTR in their monomeric and oligomeric states (Fig. 5a-c). For DRYG-CTR_P332C-C457_, increasing concentration of the unlabeled protein resulted in a second peak appearing at lower transfer efficiency. This new peak corresponds to single labeled DRYG-CTR molecules that associate to unlabeled oligomers. The fraction of this peak increases at the expense of the higher transfer efficiency peak, with increasing concentration of unlabeled protein (Fig. 5a). This behavior indicates an increase of the fraction of oligomer-bound labeled protein (Fig. 5d), in parallel with increasing fraction of oligomers (Fig. 4c). For DRYG-CTR_C457-G523C_ and DRYG-CTR_G523C-Y609C_, we instead observed an apparent gradual shift of the single peak towards lower transfer efficiencies (together with increasing the peak width at intermediate protein concentrations) (Fig. 5b,c,e). Thus, all three protein regions display a consistent reduction of mean transfer efficiency of the protein when bound to oligomers in keeping with an oligomerization-driven increase of inter-residue distance throughout the entire IDR. We wondered if the observed distance increase originates from the overall expansion of the disordered ensemble, or if it can be explained by the formation of local structure. An answer is provided by the analysis of the fluorescence lifetimes of three DRYG-CTR constructs, which systematically cluster at the dynamic line, throughout the course of oligomerization, indicating that the protein remains disordered also in the context of the oligomers (Fig. 5a-c). The same conclusion is also supported by essentially identical CD spectra at low (monomeric) and high (oligomeric) protein concentrations (Supplementary Fig. 6). Interestingly, the three regions nevertheless show variable response to oligomerization, as evidenced by different extent of mean transfer efficiency change for monomeric and oligomeric protein states (Fig. 5a-c). The DRYG-CTR_P332C-C457_ construct undergoes larger expansion (Δ*R*_DA_ *∼* 2 nm), whereas DRYG-CTR_C457-G523C_ and DRYG-CTR_G523C-Y609C_ show more modest expansion (Δ*R*_DA_ *<* 1 nm) (Supplementary Fig. 7). Thus, the protein regions at immediate vicinity to the oligomerization core (such as the DRYG-rich region) undergo larger conformational changes upon protein self-assembly, compared to the rest of the chain that senses the oligomerization to a lesser degree.

**Fig. 5:**
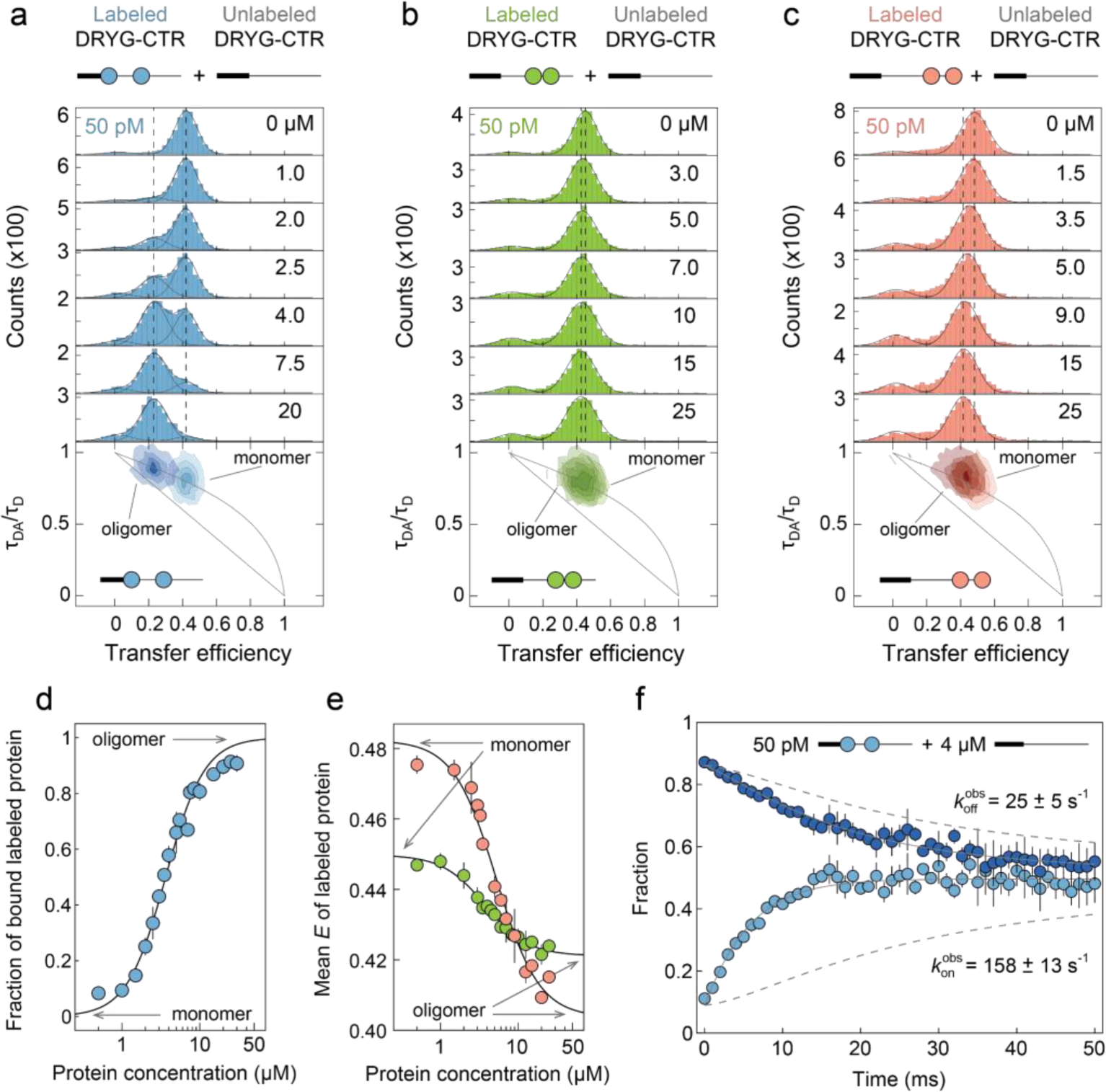
Oligomerization of eIF4B IDR and underlying conformational dynamics. **a-c** Transfer efficiency histograms of DRYG-CTR_P332C-C457_, DRYG-CTR_C457-G523C_ and DRYG-CTR_G523C-Y609C_ constructs at increasing concentrations of unlabeled protein. The dashed lines indicate mean transfer efficiencies of unbound DRYG-CTR (monomeric) and bound DRYG-CTR (oligomeric), for each construct, respectively. The bottom panels show the normalized donor lifetime - transfer efficiency histograms for each construct in the absence and presence of unlabeled protein, corresponding to monomeric and oligomeric states, respectively. The diagonal and curved lines indicate the static and dynamic lines, as in Fig. 3. **d** The fraction of oligomer-bound DRYG-CTR_P332C-C457_ as a function of overall protein concentration. **e** The mean transfer efficiencies of DRYG-CTR_C457-G523C_ and DRYG-CTR_G523C-Y609C_ as a function of overall protein concentration. The black lines are fits to Hill equation. **f** Plot of DRYG-CTR association kinetics, showing the relaxation of bound (dark blue) and unbound (light blue) states over time. The solid lines show global fits with single relaxation rates. Dashed lines show the kinetic behavior in the absence of protein association (i.e. originating from the occurrence of new molecules only).

To probe the self-association kinetics we focused on the DRYG-CTR_P332C-C457_ construct, which has sufficient separation of FRET states corresponding to the oligomer bound and unbound protein, and is thus amenable to recurrence analysis of single particles (RASP)^44^. Indeed, RASP experiments confirm the relatively fast kinetics at a ms timescale (Fig. 5f). Considering the observed on/off rates, we calculated the exchange rate between the bound and unbound proteins, *k*_ex_ ≈ 184 ± 18 s^-1^, which indicates that a single DRYG-CTR molecule on average spends ∼5 ms as part of oligomers. Thus, combining our structural and dynamical characterization we can describe the DRYG-CTR oligomers as large disordered complexes that dynamically form on the basis of several protein molecules.

### Molecular simulations confirm DRYG-mediated oligomerization and associated IDR expansion

Coarse-grained (CG) simulations have been consistently leveraged against smFRET data, and have yielded experimentally sound ensembles for a series of IDPs^45–48^. We employed CG Langevin dynamics simulations to reconstruct the ensemble of eIF4B IDR constructs (DRYG-CTR and CTR), by quantitatively matching experimental and computed mean transfer efficiencies simultaneously, across multiple eIF4B positions. To further enhance the mechanistic understanding of the oligomerization behavior of eIF4B we simulated the eIF4B IDR both as a single chain and oligomers (see Methods for details). This approach allowed us to pinpoint the role of the DRYG region in oligomerization of eIF4B, to assess the protein conformational ensemble in the oligomeric state, and to understand how the size of eIF4B oligomers responds to changes in ionic strength, as investigated by experiments. Computed mean transfer efficiencies show an excellent agreement (ρ_c_ of ∼0.8) with experimental transfer efficiencies of single chains for both the DRYG-CTR and CTR constructs (Fig. 6a-c). Simulations also recapitulate the drop in the mean transfer efficiencies observed for entirety of the CTR construct when the DRYG-rich region is truncated (Fig. 6b), with the largest effect seen for the 332-457 region. We note that for two regions (361-407 and 457-523) simulations overestimate mean FRET efficiencies by approximately 10-15%. Notably, the 361-407 region brackets the transient helical motif identified in NMR experiments (Supplementary Fig. 1), and the proline-rich 457-523 region has a high prediction of PPII helical propensity (Fig. 1b), providing a rationale of more expanded conformations. These regions most prominently contribute to the inability of simulations to account for approximately 20% (1-ρ_c_) of the variance between experimental and computed FRET efficiencies. Generally, CG potentials still lack the ability to simulate secondary structure elements in disordered chains. The model we employ only contains bonded terms for two subsequently linked beads, without taking into consideration angle and dihedral torsional terms, which contribute to the local and likely transient secondary structure propensity of IDRs.

Given the good agreement with experiments for the mean transfer efficiencies of single chains, we attempted the sampling of the eIF4B ensemble in an oligomeric state. As before, the simulated DRYG-CTR constructs within oligomers show an excellent agreement with experimental transfer efficiencies (Fig. 6d). The simulations capture the reduction of mean transfer efficiencies upon oligomerization, indicative of expansion of the conformational ensemble upon monomer-oligomer transition as observed in experiments (Fig. 5). In this case, the ρ_c_ is higher than in the case of single chain simulations (ρ_c_ = 0.9) (Fig. 6e). This result is somewhat unsurprising considering the higher level of constraints in the oligomeric state, which possibly accounts for the rigidity otherwise mediated by proline-rich regions (Fig. 1c) in the single chains. Additionally, we extracted the end-to-end distances (*R*_ee_) for the different regions labeled in smFRET experiments both for single chains and oligomers. The comparison reports on an increase of average *R*_ee_ to various degrees for three constructs, in line with the expansion of the CTR upon oligomerization, as observed in smFRET experiments (Supplementary Fig. 7). Notably, in parallel with the increase of average *R*_ee_s, the *R*_ee_ distributions get systematically narrower. Simulations, together with smFRET experiments (Fig. 5), suggest that the conformational ensemble of eIF4B CTR undergoes pronounced reshaping upon oligomerization. We then further analyzed the molecular components of the ensemble retrieved in the oligomeric state, to understand if specific regions within eIF4B IDR drive association. We quantified the map of inter-chain contacts within oligomers (see Methods for details), which shows a significantly higher fraction of contacts concentrating in the DRYG-rich region (Fig. 6f) and gradually fading towards the C-terminal end of the chain (Fig. 6f). Notably, both these observations are in line with the experimental data (Fig. 2f,g, ) identifying the DRYG-rich region as a key mediator of the oligomerization process. Moreover, the combination of experiments (NMR and smFRET) and simulations provide a highly consistent picture rationalizing the molecular principles of eIF4B IDR oligomer formation.

**Fig. 6:**
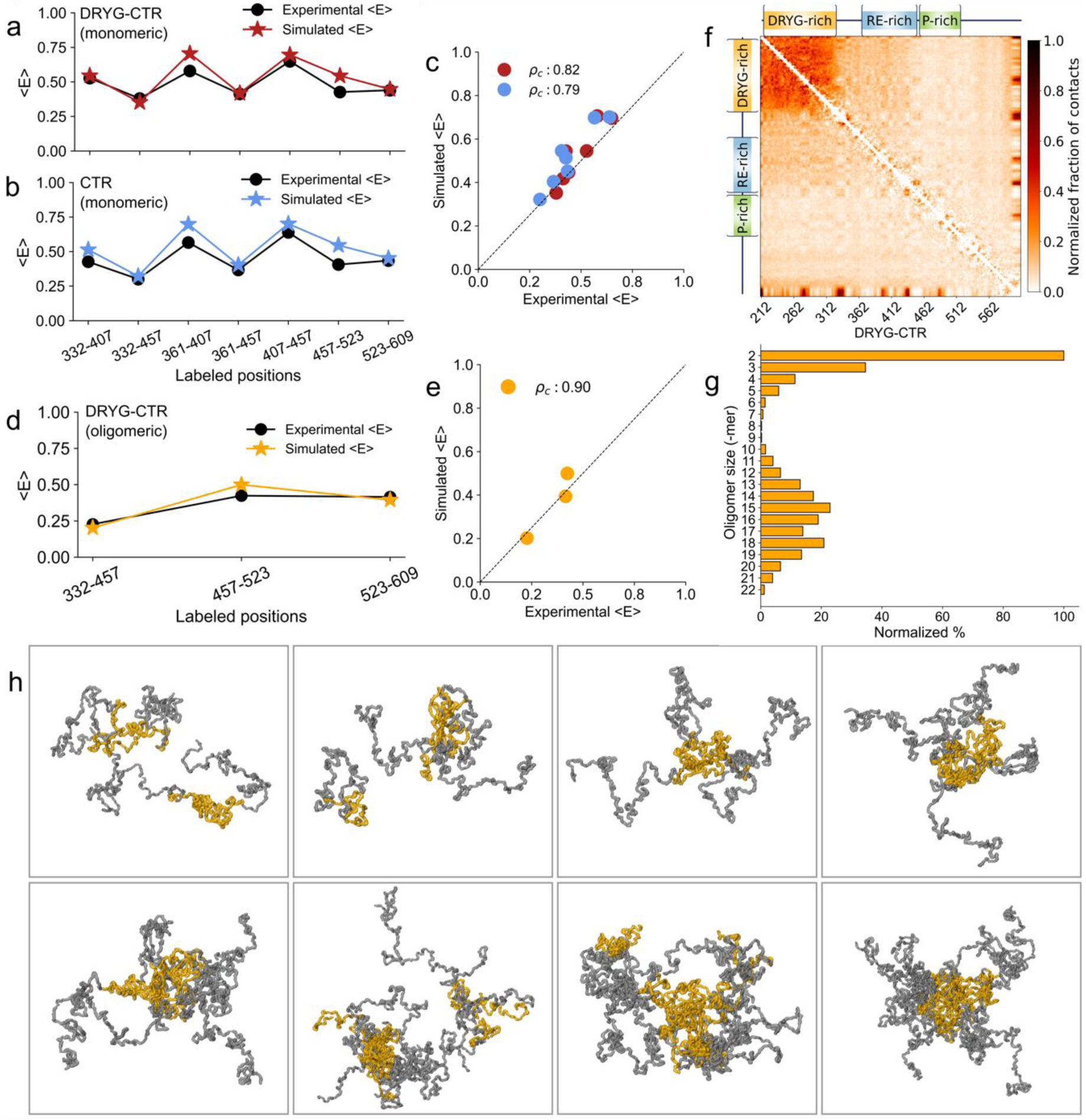
Coarse-grained ensembles of eIF4B IDR suggest direct role of the DRYG-rich region in driving oligomer formation. **a, b** Experimental (black) and simulated (red and blue) mean transfer efficiencies, 〈*E*〉 as a function of labeled positions for monomeric DRYG-CTR (a) and CTR (b) at an ionic strength of *I* = 192 mM (corresponding to buffer ionic strength of 20 mM NaP with 150 mM NaCl). **c** Correlations between the experimental and simulated 〈*E*〉. The concordance correlation coefficients (*ρ*_c_) are reported in the legend for DRYG-CTR and CTR, respectively. **d** Experimental (black) and simulated (red and blue) 〈*E*〉 as a function of labeled positions for DRYG-CTR simulated at an ionic strength of *I* = 117 mM (corresponding to buffer ionic strength of 20 mM NaP with 75 mM NaCl) and conditions favoring formation of oligomers (see Methods for details). **e** Correlation between the experimental and simulated 〈*E*〉. **f** Difference in the fraction of contacts between monomeric and oligomeric DRYG-CTR,

### Heterogenous size distribution of DRYG-CTR oligomers is sensitive to the ionic strength

As demonstrated by FCS experiments, DRYG-CTR forms large oligomers containing on average more than 10 molecules, with the average oligomer size showing a slight dependence on ionic strength. Nevertheless, these experiments do not provide information about the heterogeneity of the oligomer population or their size distribution. As a goal to expand this observation, we analyzed the populations of simulated DRYG-CTR oligomer ensembles. The results indicate a heterogenous distribution of the oligomers sizes, ranging from minimal dimers up to large oligomers (Fig. 6g). The formation of such heterogeneous size oligomer population (Fig. 6h), is in excellent agreement with several experimental observations, such as the monotonous shift of FCS curves with increasing oligomer fraction (Fig. 4b), as opposed to a simple two-state behavior expected in case of monomer to fixed-size oligomer transition, as well as the apparent positive cooperativity in monomer-oligomer transition curves (that is binding isotherm with n>1) (Fig. 4c, 5d,e). To compare with the experiments, we also looked at the sensitivity of oligomerization to changes in ionic strength and analyzed the oligomer size distributions at low and high salt conditions (Supplementary fig. 11). Interestingly, at low ionic strengths of *I* = 92 and 117 mM (corresponding to buffer ionic strengths of 20 mM NaP with 50 and 75 mM NaCl, respectively) the oligomer size distribution appears bimodal. One fraction of the ensemble is populated by smaller oligomers (2-mers to 4-mers), while another fraction is based on larger sized oligomers comprising 10 and more protein molecules (Fig. 6g and Supplementary fig. 11,a,b). Comparing these two conditions, the lower ionic strength favors the formation of larger oligomers, as evidenced by an increase of their population at the expense of the smaller oligomers and a pronounced increase in the overall size of the larger oligomers. Increasing the ionic strengths to *I* = 192 and 342 mM (corresponding to buffer ionic strengths of 20 mM NaP with 150 and 300 mM NaCl, respectively), results in an appearance of tailed distributions. Here the majority of oligomers are observed in the range of 2 - 4-mers, followed by gradually decreasing smaller fraction of larger oligomers (up to ∼50-mers) (Supplementary fig. 11c,d). The formation of a defined population of large-sized oligomers at low ionic strengths is likely to be dictated by a deficient electrostatic screening, which, additionally favors protein association into larger complexes. Based on the observed oligomer size distributions, the average oligomer size appears to be dependent on the ionic strength, systematically decreasing from approximately 16-mers to 8-mers as the ionic strength increases within *I* = 92 – 342 mM. This trend is in excellent agreement with the experimental results (Fig. 4c), moreover, the range of average oligomer sizes observed in simulations is also in line with our estimates from the FCS experiments (Fig. 4c). Thus, in line with weak ionic strength dependence observed in DRYG-CTR oligomerization experiments (Fig. 4c,d), the size and distribution of DRYG-CTR oligomers vary with changes in salt concentration.

### The interplay between DRYG and CTR determines the eIF4B IDR self-association behavior

As we noted earlier, upon saturation of DRYG-CTR, the protein self-association culminates in mesoscopic phase separation and formation of condensed protein-rich droplets (Fig. 2a-c). Already through our initial qualitative characterization we noted that DRYG-rich region has extremely high phase separation propensity, which however decreases in the presence of the CTR. To rationalize this observation, we quantified the phase separation propensity of the isolated DRYG or CTR segments of the protein (Fig. 7a) and under the same conditions as the DRYG-CTR (Fig. 2c). The comparison of saturation concentrations for these variants (Fig. 7b), indicates that the DRYG-rich region is indeed essential for the phase separation of the eIF4B IDR. In the absence of this region the CTR construct does not show any detectable phase separation under physiological protein concentration and ionic strength, whereas the DRYG alone largely phase separates at very low protein concentrations even in solutions of high ionic strength (*c*_sat_∼0.2 μM in 1 M NaCl). Interestingly, the extremely low phase separation threshold of DRYG is increased by an order of magnitude in the presence of CTR, as evidenced by 2-3 μM saturation concentration of DRYG-CTR. This indicates that the CTR acts as a phase separation dampener, lowering the extremely strong intrinsic phase separation propensity of DRYG-rich region. To dissect the role of CTR in inhibition of DRYG-driven phase separation we designed two additional deletion constructs (DRYG-CTR-N and DRYG-CTR-C), in which the C-terminal or N-terminal halves of the CTR were removed, respectively (Fig. 7a). The CTR-N segment encompasses the entire RE-rich region, almost evenly interspersed by positively and negatively charged residues (Fig. 1c), and is slightly acidic at neutral pH (net charge is -1). The CTR-C segment on the other hand encompasses the entire P-rich region and the heavily charged far C-terminal region and has a net charge of -5, particularly due to a cluster of negatively charged residues at the very end of the protein (Fig. 1c). Interestingly, the CTR-N has relatively modest inhibitory effect on DRYG-driven phase separation and increases the saturation concentration of DRYG by less than a factor of 5 (Fig. 7b). The effect of CTR-C is much more dramatic as it raises the phase separation threshold by several orders of magnitude. This could be correlated with its acidic nature, suggesting that the overall net charge of the segments downstream of DRYG-rich region defines the saturation concentration of the protein, as also observed with other IDRs^49–51^.

To rationalize the mechanisms of DRYG-CTR self-association and identify the link between the phase separation and oligomerization we quantified the phase separation saturation concentrations at different ionic strengths (Fig. 7c) and compared them with oligomerization affinities also acquired at varying salt concentrations. Interestingly, the ionic strength has a pronounced effect on DRYG-CTR phase separation propensity, as seen from the large increase of the saturation concentrations at high salt concentrations. This behavior is in striking contrast to the weak ionic strength dependence of DRYG-CTR oligomerization (Fig. 4c,d and 7c), thus indicating that additional electrostatically-driven molecular mechanisms are behind the phase separation process. Notably, the ionic strength dependence of oligomerization affinity and phase separation propensity for DRYG-CTR-N and DRYG-CTR-C constructs follow similar trends as for DRYG-CTR (Fig. 7c). Depending on the actual CTR sequence, the ionic strength dependent phase diagram (i.e. *c*_sat_ values) and oligomerization affinity (*K*_D_ values) laterally shift along the diagonal of protein - NaCl concentration plane (Fig. 7c). The comparison of these three constructs thus points towards a pronounced coupling between the oligomerization and phase separation processes. Thus, the DRYG-DRYG mediated interaction is an essential step for self-association of eIF4B IDR, whereas the CTR acts as a regulator of self-association efficacy. The balance between these two features defines both the self-association affinity and the condensation threshold at the two extremes of eIF4B IDR self-association landscape, such as oligomerization and phase separation.

**Fig. 7:**
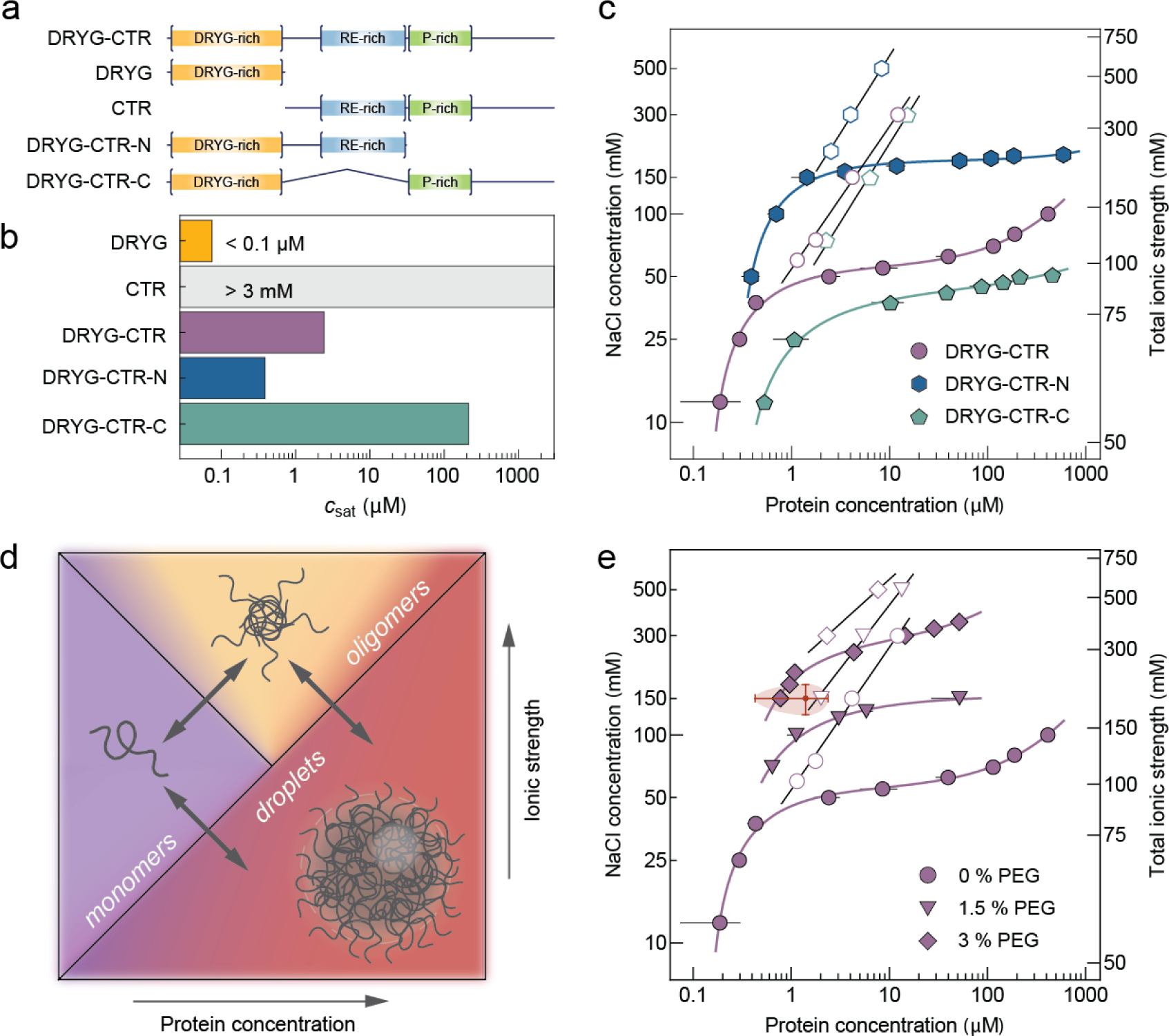
Disordered segments of eIF4B define its phase separation behavior. **a** Schematics of DRYG-CTR and truncated constructs used in phase separation experiments. **b** Bar chart showing the *c*_sat_ values for DRYG-CTR and truncated constructs measured at 20 mM NaP, 50 mM NaCl (pH 7.0) at 20 °C. **c** Phase diagrams of DRYG-CTR, DRYG-CTR-N and DRYG-CTR-C constructs based on *c*_sat_ values (filled circles) and *K*_D_ values (open symbols, as in Fig. 4c and Supplementary Fig. 12) as a function of NaCl concentration and total ionic strength (right axis). The solid colored lines are fits to an empirical dose-response function. The solid black lines are fits with the Lohman–Record model^43^. **d** Schematic phase diagram defined by protein concentration and ionic strength indicative of the self-association landscape of eIF4B IDR and underlying transitions across monomeric, oligomeric and condensed droplet states. **e** Phase diagrams of the DRYG-CTR construct based on *c*_sat_ values (filled circles) and *K*_D_ values (open symbols) as a function of NaCl concentration and total ionic strength (right axis), in the absence and presence of crowding agent PEG. The solid colored lines are fits to an empirical dose-response function. The solid black lines are fits with the Lohman–Record model^43^. The red point indicates the average eIF4B concentration in HeLa cells^52–54^ and a range of physiological ionic strength. The shaded area shows the mean deviation along both axes.

## Discussion

Human translation initiation factor eIF4B remains one of the least structurally and functionally characterized translation factors. In contrast to the human protein, the yeast ortholog of eIF4B was studied more extensively. Rationalizing the structural and functional behavior of human eIF4B based on these data, is however, challenging, as the two orthologues share minimal sequence conservation (∼20 %) and are not fully equivalent in function^55^. One of the unique features of the human homolog is the presence of the DRYG-rich region within the long IDR, which was shown to promote eIF4B self-association^32^. Notably the yeast ortholog lacks a DRYG-like region and its self-association was not reported. In this work we integrated a large set of experiments and simulations to structurally characterize the human eIF4B IDR in its various molecular states across different regimes of the self-association landscape. We have described the conformational ensemble of complete eIF4B IDR in its monomeric form and were able to detect and describe the distinct population of large and dynamically forming oligomers. Importantly, our integrative approach enabled us not only to describe the intricate details of oligomerization, but also follow the conformational reshaping of the disordered ensemble across the monomer - oligomer transition and quantify the underlying dynamics at intra- and intermolecular scales.

The growing evidence indicates that for many condensation-prone proteins, the propensity of *in vitro* phase separation, as well as formation of biological condensates, is often correlated with initial formation of large-size oligomers/clusters at concentrations far below the condensation thresholds^49,56–60^. Here we show that the *in vitro* self-association landscape of eIF4B follows a similar scenario, as exemplified by the behavior of its key self-associating long disordered region. The phase diagram of eIF4B IDR shows three distinct regions across the two-dimensional space of protein concentration *versus* ionic strength, corresponding to monomeric, oligomeric and condensed (droplets) states (Fig. 7d). Although a narrow range of low ionic strengths strongly favors a direct phase transition from dispersed monomeric phase, at higher and physiologically relevant ionic strengths the formation of large-size oligomers precedes the phase separation and appear to be a critical and mandatory step along the condensation pathway (Fig. 7). Notably, DRYG-rich region is at the core of both oligomerization and phase separation, as neither the N-terminal RRM domain (and its tail)^32^ nor the C-terminal IDR (i.e. the CTR; Fig. 7b) that flank the DRYG-rich region, exhibit any detectable self-association properties.

Although DRYG-DRYG association is essential for self-association at nanoscale (i.e. oligomer formation) and mesoscale (i.e. droplet formation), the two processes demonstrate differing responses to ionic strength (Fig. 7c). The weak ionic strength dependent oligomerization we attribute to the intermolecular cation-π and π-π interactions, likely mediated by arginine-tyrosine and tyrosine-tyrosine interactions, similar to the intramolecular contacts that define the compact conformational ensemble for monomeric DRYG (Fig. 3). In contrast, the pronounced ionic strength dependence suggests an electrostatic driving force behind the phase separation, likely orchestrated through interactions between aspartate and arginine residues of the DRYG-rich region. These interactions are favored at low ionic strength regime with poor electrostatic screening, leading to rapid coalescence of monomers into phase separated droplets. Higher ionic strengths weaken these contacts and raise the phase separation threshold. However, the formation of oligomers amplifies the effective interaction strength by increasing the interaction valency as a function of oligomer size. Not surprising, the phase separation propensity is also correlated with the fraction of larger size oligomers (Fig. 4c and Supplementary Fig. 11, 12). Therefore, given the smaller oligomers typically observed at high salt concentrations a higher overall protein concentration will be required to nucleate eIF4B IDR condensation and to compensate for the penalty of lower interaction valency per oligomer.

While the DRYG-rich region orchestrates the self-association of eIF4B, the CTR defines and fine-tunes the precise boundaries across the self-association landscape. Comparing DRYG-CTR constructs with varying CTR lengths demonstrates that the effect of CTR, likely mediated by charged interactions, is strongly correlated between the protein oligomerization and phase separation. This property, thus systematically couples the eIF4B oligomerization affinity with phase separation propensity. This is an interesting observation, as it identifies the CTR as a point of control over eIF4B self-association at a global scale. Interestingly, the sequence conservation analysis assigns high scores to several polypeptide segments within the CTR (Supplementary Fig. 13), which coincide well with the known phosphorylation and previously mapped interaction sites^9,11,30,32^. Phosphorylation is a well-known modulator of protein phase separation^61–63^, thus it is reasonable to expect that eIF4B phosphorylation at previously identified functionally relevant Ser406 and Ser422 sites^22^ located in the CTR, may have a strong impact on protein self-association behavior. Similar, self-association regulation could also be achieved by means of reentrant phase transition, for example upon RNA binding to CTR. The potential of such regulation is further supported by the observation that phase separation behavior of the eIF4B IDR is extremely sensitive *in vitro*. In addition to the pronounced effect of ionic strength on eIF4B self-association (Fig. 7c), the phase behavior is also very sensitive to molecular crowding (Fig. 7e). Even minimal crowding introduced by PEG strongly influences the phase diagram, counteracting the effect of ionic strength, and shifting the phase diagram towards a more physiologically relevant ionic strength regime. Notably, under these conditions the “triple point” of the phase diagram is positioned at the close proximity to the estimated cellular eIF4B concentration and physiological ionic strength (Fig. 7e). Although beyond the scope of this work, such behavior supports our expectation that the monomer-oligomer-condensate transitions that we observe *in vitro* might also be relevant at the cellular level and be used as a response and coping mechanism to rapid changes of cellular microenvironment under stress condition, such as variation of the intracellular ionic strength or crowding upon osmotic stress^64–67^. In this context, *in-cell* smFRET experiments^68^ might open up future possibilities for monitoring self-association transitions (e.g. monomer-oligomer transitions) directly in live cells.

It remains to be determined how oligomerization of eIF4B is utilized functionally during translation initiation and stress granule formation. Beyond previously proposed hypotheses, such as eIF4B-dimerization dependent RNA annealing activity^55^ or inhibition of interaction with eIF3^32^, the formation of large “fuzzy” oligomers may be a sensitive and dynamic mechanism to enhance the effective binding affinity of the protein towards its targets, such as mRNA. Moreover, the pronounced reshaping and systematic expansion of the conformational ensemble of eIF4B IDR upon oligomerization (Fig. 5) may further enable oligomerization-driven tuning of the accessibility and affinity towards its binding partners. Our work thus provides detailed structural insights beyond simple dimerization of eIF4B, which can serve as a basis for future validation of these important hypotheses.

## Methods

### Protein expression and purification

A synthetic gene based on the protein sequence of the human eIF4B protein (Uniprot entry P23588) was purchased as codon-optimized DNA for *E. coli* expression (Supplementary table 1) (Integrated DNA Technologies). All constructs were generated based on PCR amplification of specific regions, and inserted into the *Ssp*I site of the 2B-T modified pET vector (Addgene plasmid #29666) using the ligation-independent cloning protocol. The vector also adds an N-terminal hexahistidine (His_6_) tag followed by a tobacco etch virus (TEV) protease cleavage site on the expressed proteins. PCR-based site-directed mutagenesis was performed to create double site-specific cysteine constructs and truncated variants (see Supplementary table 2 for the list of primers used). The full list of constructs is presented in Supplementary table 3.

For protein expression, colonies of freshly transformed *E. coli* T7 Express *lysY* (New England Biolabs) were first grown overnight in 10 mL of lysogeny broth (LB) with 50 µg/ml ampicillin, then transferred to 1 L cultures at 37 °C in TB or 500 ml in M9 minimal media containing 1 g/L [^15^N]NH_4_Cl. At an OD_600nm_ = 1.8 for TB or OD_600nm_ = 0.6 for M9, protein expression was induced with a final concentration of 0.5 mM isopropyl β-D-1-thiogalactopyranoside (IPTG) followed by overnight growth at 20 °C. Bacteria were harvested by centrifugation and then resuspended in a lysis buffer consisting of 20 mM Tris-HCl (pH 7.8), 250 mM KCl, 20 mM imidazole, 100 mg/L lysozyme, 1 mM PMSF, 2 mM β-mercaptoethanol, 0.5-2 M of urea (depending on the construct) and one protease inhibitor cocktail tablet (Roche). Following sonication on ice (10 min at 40 % power, alternating 50 s of sonication and 60s of pause), the sample was centrifuged at 40000 x *g* for 40 min at 4 °C to remove cellular debris. Specifically, for the DRYG-CTR-N variant, the bacteria were harvested by centrifugation and then resuspended in a Triton-containing lysis buffer consisting of 50 mM Tris-HCl (pH 7.8), 250 mM KCl, 1% Triton-X100, 2 mM β-mercaptoethanol, 1 M of urea. Following sonication on ice (1 min at 40% power, alternating 20s of sonication and 20s of pause), the sample was centrifuged at 20000 x *g* for 20 min at 4 °C to remove cellular debris. The supernatant was filtered with a 0.7 µm glass Microfilter GF/F (GE Healthcare Life Sciences Whatman) and added to NUVIA Ni^2+^ affinity chromatography resin (Bio-Rad) in a plastic column, washed first with five column volumes of binding buffer (20 mM Tris-HCl (pH 7.8), 250 mM KCl, 20 mM imidazole, 2 mM β-mercaptoethanol), then three column volumes with a higher salt concentration (800 mM of KCl), and finally five column volumes with increased imidazole concentration (30 mM). The protein was eluted with a buffer containing 20 mM Tris-HCl (pH 7.8), 250 mM KCl, 500 mM imidazole, 2 mM β-mercaptoethanol. The cleavage of the N-terminal His-tag for DRYG-CTR constructs was performed by TEV protease (15 µg/mg protein) during a dialysis step against 20 mM Tris-HCl (pH 7.8), 250 mM KCl, 5 mM β-mercaptoethanol with 0.5M urea and 1% glycerol. The same procedure for CTR constructs was performed in 20 mM Tris-HCl (pH 7.8), 250 mM KCl, 5 mM β-mercaptoethanol buffer. The protease, His-tag and remaining uncleaved protein were removed by a second Ni^2+^ affinity chromatography step, followed by a purification with anion exchange chromatography with EnrichQ (Bio-rad), MonoQ, HiTrap Q or HiTrap DEAE (Cytiva) columns. In addition, prior to fluorescent labeling, protein samples were reduced by TCEP and were further purified with reverse-phase HPLC on a C18 column (ReproSil Gold 200, Dr. Maisch), followed by lyophilization, yielding samples of 2-5 mg dried protein depending on the size of the column used. Protein samples for NMR were used after anion exchange purification and were concentrated and buffer exchanged into corresponding NMR buffers.

### Fluorescent labeling of proteins

For single-molecule FRET experiments, double cysteine constructs of proteins were fluorescently labeled with Cy3B (Cytiva) and CF660R (Biotium), as donor and acceptor dyes, respectively. The labeling was performed under denaturing conditions (100 mM NaP, 6 M GdmCl, pH 7.1), in two steps. First, the protein was incubated with the Cy3B maleimide at a protein to dye ratio 1:0.7, for 1 h at room temperature, followed by purification of the labeling mixture with reverse-phase HPLC, on a C18 column (ReproSil Gold 200, Dr. Maisch). This procedure enabled separation of unlabeled, single-labeled (in most of the cases as two separate single site-specifically labeled protein permutants) and double-labeled protein fractions, as well as the small fraction of unreacted/inactive dye. All corresponding protein fractions were collected individually and were lyophilized overnight. In a second step, one of the single labeled protein fractions (corresponding to a particular site-specifically donor-labeled protein) was further labeled with CF660R maleimide at a protein to dye ratio 1:3, for 2-3 h at room temperature, followed by purification of the labeling mixture with reverse-phase HPLC, on a C18 column (ReproSil Gold 200, Dr. Maisch). This procedure enabled the separation of the single-labeled and double-labeled protein fractions, as well as the excess of the unreacted acceptor dye. After these rigorous labeling and purification steps, it was possible to obtain predominantly double labeled protein, in most cases with site-specific attachment of donor and acceptor dyes. For FCS experiments, the proteins were single-labeled with ATTO 655 maleimide dye and purified using a similar procedure as described above.

### Phase separation assays

For phase separation experiments the lyophilized protein was dissolved in 20 mM NaP, 1mM TCEP, pH 7.0 buffer containing 100-300 mM NaCl. The dissolved protein solution was then repeatedly buffer exchanged with a centrifugal filter (Amicon, Merck) against the same buffer to eliminate the residual TFA, remaining after reverse-phase HPLC and lyophilization and was concentrated up to ∼1mM, to yield concentrated one-phase protein solution. The DRYG construct was dissolved and concentrated in double-distilled water and was used as it is.

The light phase or saturation concentrations, *c*_sat_, as well as the dense phase concentrations, *c*_dense_, were obtained by the centrifugation method^69^. In brief, at respective NaCl concentration, a protein solution was prepared under conditions favoring phase separation, which was confirmed by solution turbidity and observation of droplets under the microscope. The phase separated solution was then incubated at respective temperatures in a thermocycler (Mastercycler nexus, Eppendorf) for 20 minutes, followed by centrifugation at 20500x g for 5 min at respective temperature, using a temperature-controlled centrifuge (Eppendorf 5430R or 5425R). After 5 minutes, the dense phase is settled as a pellet and the light phase remains as supernatant. A portion of supernatant was collected without disturbing the pellet and was diluted in 6 M GdmCl. Similarly, 1μl of dense phase was collected using a positive displacement pipette (Microman E, Gilson) and was also diluted in 6 M GdmCl. The protein concentrations were measured spectrophotometrically (Cary 60, Agilent Technologies) with 1 mm pathlength tray cell (Hellma) or 1 cm quartz cuvettes (Hellma). The *c*_sat_ and *c*_dense_ values were then estimated with consideration of the dilution factors. Each data point was repeated at least three times. The *c*_sat_ and *c*_dense_ values were found to be independent of the initial volumes of the two-phase solution as well as the centrifugation time and speed. For phase separation assays in the presence of a crowding agent, PEG 8000 (P1458, Sigma-Aldrich) was used.

### Differential interference contrast (DIC) and fluorescence imaging

DIC images were acquired using an IX83 (Olympus) brightfield microscope equipped with CMOS camera (Thorlabs), polarizer (IX-LWPO, Olympus), analyzer (IX3-AN, Olympus), Nomarski prism (IX2-DIC60, Olympus) and a water immersion objective (UplanSApo 60x/1.20W, Olympus). The fluorescence images were acquired using a MicroTime 200 confocal microscope (PicoQuant), employing a 520 nm laser (LDH-D-C-520; PicoQuant) operating in a continuous-wave mode and a XY piezo scanning stage (PI). The solutions with phase separated DRYG-CTR droplets were prepared at 5 μM protein concentration in 20 mM NaP (pH 7.0), 50 mM NaCl, 1 mM TCEP and were imaged in ibidi multiwell chambers (µ-Slide 15 Well 3D, ibidi). For fluorescence imaging the solution was doped with 20 nM of Cy3B-labeled DRYG-CTR.

### CD spectroscopy

The CD spectra were recorded with a Jasco J-1500 CD spectrometer equipped with a temperature-controlled cuvette holder, using 0.1 or 1 mm path-length cuvettes at 20 °C. The CD spectra of DRYG-CTR were measured in a buffer containing 20 mM NaP (pH 7.0), 75 mM NaCl or 300 mM NaF, 1 mM TCEP. The CD spectra of CTR, CTR-N, CTR-C were measured in a buffer containing 20 mM NaP (pH 7.0), 150 mM NaF, 1 mM TCEP. All CD spectra were recorded between 190 and 250 nm, with data pitch 1 nm, scan speed 50 nm/min and accumulation of 10 scans. Before recording the spectra, baseline correction was performed with the respective buffers.

### NMR spectroscopy

NMR data were collected on a Bruker Avance 700 or 800 MHz spectrometer, equipped with triple-resonance gradient room-temperature or cryogenic probe, respectively. NMR data were processed by using NMRPipe/NMRDraw software^70^ and NMR spectra were analyzed using Sparky (T. D. Goddard & D. G. Kneller, University of California, San Francisco, USA). Samples contained 170 μL in a 3mm NMR tube at 293 K, in a buffer containing 20 mM NaP (pH 7.0), 2 mM DTT, and varying amounts of NaCl and GdmCl (deuterated). Chemical shift assignments for eIF4B-CTR were previously deposited in the BMRB (https://bmrb.io/) under accession codes 51952, 51953 and 51957^29^.

### Single-molecule fluorescence spectroscopy

Single-molecule fluorescence measurements were performed using a MicroTime 200 confocal microscope (PicoQuant). Alternating excitation of the dyes was achieved using pulsed interleaved excitation (PIE)^71^, by synchronizing 520 nm (LDH-D-C-520; PicoQuant) and 638 nm (LDH-D-C-640; PicoQuant) diode lasers at 20 MHz pulse rate with a delay time of 25 ns between the two laser pulses. The power of 520 nm laser excitation was adjusted to 100 µW (measured at the back aperture of the microscope objective), whereas the direct acceptor 638 nm excitation was adjusted to ∼50 μW (such that the donor and acceptor emission intensities were matching after donor and direct acceptor excitations, respectively). The lasers were focused on the sample through a water immersion objective (UplanSApo 60x/1.20W, Olympus), 50 μm in the solution above the cover slide surface. The emitted photons from the sample were collected by the same objective, and were separated from the excitation photons and scattered light by a major dichroic mirror (ZT532/640rpc, Chroma) and 100 μm pinhole (Thorlabs), respectively. The remaining photons were then detected onto four detection channels (two donor and two acceptor) based on their polarization (using a polarizing beam splitter) and wavelength by two dichroic mirrors (T635lpxr, Chroma). The donor and acceptor photons were then further filtered by bandpass filters (ET585/65 and H690/70, Chroma) before being focused onto the SPAD detectors (SPCM AQRH-14-TR, Excelitas Technologies). The arrival time of each photon was recorded with a HydraHarp 400 TCSPC module (PicoQuant) with 16 ps time resolution.

All smFRET measurements were performed with 50-100 pM of double labeled protein solution in 20 mM NaP (pH 7.0) buffer at corresponding concentrations of NaCl or GdmCl, supplemented with 0.01% Tween 20 and 143 mM β-mercaptoethanol. The smFRET self-association assays were performed under the same solution conditions, but with increasing concentrations of unlabeled proteins. Each sample was measured at room temperature (20 ± 1°C) with acquisition time of 15-30 mins, each dataset was repeated at least once. All measurements of DRYG-CTR constructs were performed in ibidi multiwell chambers with uncoated polymer coverslip (µ-Slide 15 Well 3D, ibidi). For measurements of DRYG-CTR at low NaCl (<100 mM), the ibidi chambers were additionally coated with PLL-PEG (SUSOS, Switzerland). PLL-PEG coating was achieved by incubation of wells with 0.5 g/L solution of PLL-PEG for 10 min, followed by a wash with distilled water and respective measurement buffer. All measurements for CTR constructs were performed in custom-made PEG coated glass cuvettes, which were prepared according to previously reported protocol^72^.

smFRET data analysis was performed using the Fretica package for Mathematica (Wolfram Research Inc, USA) developed by Daniel Nettels and Benjamin Schuler (https://schuler.bioc.uzh.ch/programs/). Briefly, fluorescence bursts were identified by ΔT method, with interphoton time of 150 μs and a minimum number of 60-80 photons per burst. Transfer efficiencies for each burst were calculated according to:

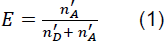

where 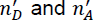 are the donor and acceptor counts (i.e. numbers of photons), respectively, corrected for background, channel crosstalk, acceptor direct excitation, differences in detector efficiencies, and quantum yields of the dyes, as previously described^73,74^. The stoichiometry ratio was calculated according to:

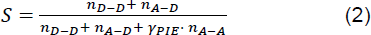

where 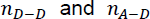 are the background corrected total donor and acceptor counts, respectively, after donor excitation (520 nm laser) and *n*_*A−A*_ is the total acceptor counts after acceptor direct excitation (638 nm laser). *γ_PIE_* is a correction factor used for weighing the *n_A−A_* to account for differences in 520 and 638 nm laser intensities. The final transfer efficiency histograms were built based on bursts originating from molecules containing both donor-acceptor fluorophores (thus eliminating molecules with photobleached acceptor or donor, i.e. donor-only and acceptor-only sub-populations). These bursts were selected based on stoichiometry ratio values 0.2 < *S* < 0.8.

To determine the mean transfer efficiencies for different sub-populations, the transfer efficiency histograms were fitted by a sum of 2-3 Gaussian peak functions. When analyzing histogram series, a global fitting was used with some fitting variables (e.g. width or position) being shared within the same histogram series. The small peak at E ≈ 0, sometimes appearing at transfer efficiency histograms, originates from the small fraction of photobleaching and blinking molecules still remaining after the PIE-cut as well as possible residual fluorescent impurities of the buffer.

### Conversion of mean transfer efficiency to mean distances

For conversion of mean transfer efficiency into mean inter-residue (donor-acceptor) distances, the following equation was numerically solved:

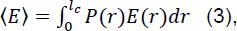

where the *l*_c_ is the contour length of the probed polypeptide chain segment, and

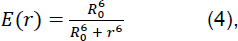

with the *R*_0_ as Förster radius at particular buffer conditions. *R*_0_ = 5.9 nm was calculated for Cy3B/CF660R dye pair, with the solution refractive index equal to 1.333, which was further scaled to account for differences in the refractive index values of particular buffers. Solution refractive index values were measured using temperature-controlled digital Abbe refractometer (DR-A1-Plus, Atago) at 20 °C.

For the *P*(*r*) distance distribution we used the SAW-ν model^75^, which adopts the following form:

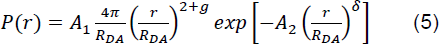

where, 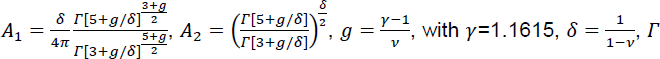 is the gamma function and ν is the scaling exponent. Eq. 5 has two adjustable parameters, the scaling exponent ν and the mean inter-residue distance, *R_DA_*, (i.e. distance between the two dyes), which equates to the root mean squared end-to-end distance, 〈*r*^2^〉^1/2^. The mean end-to-end distance for unfolded proteins approximately follows the scaling law:

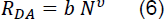

where the 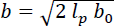 is an empirical prefactor, *l_p_* is the persistence length, *b_o_*=0.38 nm is the peptide bond length, and *N* is the number of amino acids between the two dyes (including the contribution of the dye linkers, with each approximated as *l*=4.5 peptide bond). The prefactor *b* has been previously estimated to be equal approximately 0.55 nm for proteins (with *l_p_*=0.4 nm). With this approximation, it is possible to solve the Eq. 5 for both *R_DA_* and ν.

### Nanosecond fluorescence correlation spectroscopy (nsFCS) measurements

The nsFCS measurements were carried out with the MicroTime 200 confocal microscope (PicoQuant) with the 520 nm diode laser (LDH-D-C-520; PicoQuant) operating in continuous-wave mode at a laser power of 100 μW. The emitted photons (after the pinhole) were split by a 50/50 beam splitter before being separated and filtered based on wavelength and detected at respective donor and acceptor detectors. The typical data acquisition times for nsFCS measurements were 16–20 h. All measurements were performed at protein concentration of ∼1-2 nM, in 20mM NaP (pH 7.0), 0.01 % Tween-20, 143 mM β-mercaptoethanol at respective NaCl/GdmCl concentrations. The measurements at each condition were repeated 1-2 times. The autocorrelation curves of donor and acceptor channels and cross-correlation curves between donor and acceptor channels were calculated from the measurements following the previously described methods^39,76^. The calculated correlation curves (for delay times up to 4 μs) were fitted with the following equation:

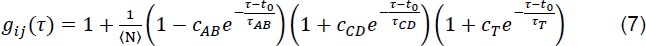

where *i* and *j* correspond to the type of signal (e.g. D or A, corresponding to the signal either from the donor or acceptor channels, respectively) and 〈N〉 is the mean number of molecules in the confocal volume. The three multiplicative terms describe the contributions of photon antibunching (AB), chain dynamics (CD), and triplet blinking (T) of the dyes, with corresponding amplitudes, *c*, and relaxation times, *τ*, respectively. *τ_CD_* was then converted in the reconfiguration time *τ*_*r*_ of the interdye segment as previously described^77^. An additional _−_ multiplicative term 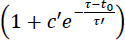 was used for D-D and A-A auto-correlations to describe the fast decay observed on a short timescale (<10ns). The timescale, *τ*′, for this decay is 6-7 ns and it can be associated with the rotational motion/tumbling of the overall protein. The same multiplicative term was also included for D-A cross-correlation for fitting the data of DRYG-CTR_P213C-P332C_ construct in the presence of NaCl. The three correlation curves from each measurement were fit globally with the time origin, t_0_, and *τ*′ as shared fit parameters.

### Fluorescence correlation spectroscopy (FCS) measurements

The FCS measurements were performed with the MicroTime 200 confocal microscope with the 638 nm diode laser (LDH-D-C-640; PicoQuant or LBX-638; Oxxius) operating in continuous-wave mode at a laser power of 50 μW. The emitted photons (after the 100 μm pinhole) were split by a 50/50 beam splitter and detected on two detectors. The typical data acquisition times for FCS measurements were 120 seconds with each sample measured 5 times and each experiment was repeated at least once. All measurements were performed at protein concentration of ∼250 pM, in 20mM NaP (pH 7.0), 0.01 % Tween-20, 143 mM β-mercaptoethanol at respective NaCl concentrations, in the absence and presence of unlabeled proteins. The correlation curves were fitted with using a three-dimensional free diffusion model (Eq. 8) with one diffusional component and an additional component for triplet blinking:

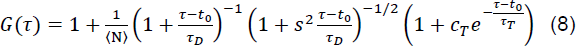

where 〈N〉 is the mean number of molecules in the confocal volume, *c_T_* is the amplitude of triplet component, *τ_D_* and *τ_T_* are the diffusion time and triplet blinking relaxation time, respectively, and *s* is the axial ratio of the confocal volume. The relative diffusion rates (*D*/*D*_0_) were calculated based on the *τ_D_* ∝ *D*^−1^ relationship.

The curves corresponding to the dependence of relative diffusion on protein concentration were fitted globally with the Hill equation, with n being shared among all datasets:

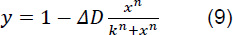

where *k* ≡ *K_D_* is the apparent dissociation constant and n is the cooperativity coefficient.

The oligomer size, shown on the right axis of Fig. 4c and Supplementary Fig. 12a,b was estimated based on 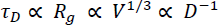 relationship, taking into account ∼15 % increase of the CTR radius of gyration upon oligomerization.

### Recurrence analysis of single particles (RASP) measurements

The RASP^44^ experiments were performed with the MicroTime 200 confocal microscope (PicoQuant) with the 520 nm (LDH-D-C-520; PicoQuant) and 640 nm (LDH-D-C-640; PicoQuant) lasers operating in PIE mode with laser powers adjusted to 100 and 50 μW, respectively. Samples consisted of 25-50 pM double-labeled DRYG-CTR with respective µM concentration of unlabeled protein, prepared in 20mM NaP (pH 7.0), 75 mM NaCl, 0.01 % Tween-20, 143 mM β-mercaptoethanol buffer. Each sample was typically acquired for ∼20 h, to ensure sufficient burst statistics. Photon bursts were first identified by combining successive photons that were separated by less than 150 µs and only bursts containing > 60 photons were retained for the further analysis. These bursts were then time-binned with a binning time of 1 ms and only the time intervals containing more than 40 photons were used for the recurrence analysis. To generate recurrence histograms first the initial transfer efficiency ranges of *E* ∈ [0.15 – 0.25] and *E* ∈ [0.4 – 0.55] were selected to define the bound and unbound populations, respectively. The burst list retained for recurrence analysis containing at least 200,000 bursts per experiment, was then scanned for burst pairs for which the first burst matched the given *E* range (either bound or unbound *E* range), followed by a second burst detected within a given time interval (i.e. recurrence interval). From the collection of second bursts, 2 sets of recurrence transfer efficiencies histograms were built corresponding to first bursts originating from either unbound or bound population, for recurrence time interval of 1 – 50 ms with a step of 1 ms. The resulting recurrence histograms were then fit with a sum of 2 Gaussian peak function to describe the unbound and bound FRET populations. The fraction of each population was calculated ratiometrically based on the area under each peak and was plotted as a function of the recurrence time. The resulting curves were fitted with the following equation:

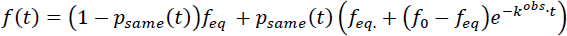

where *f*_eq_ is the transfer efficiency at equilibrium (i.e. *t* = ∞), *f*_o_ is the fraction of either bound or unbound population at *t* = 0 ms, *k^obs^* is the observed binding or unbinding rate (i.e. 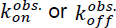, respectively) and *p_same_*(*t*) is the probability that the second burst originates from the same molecule as the first burst, which can be independently quantified from the data^44^. Each experiment was repeated three times, the resulting kinetic curves were individually fitted and the resulting rate coefficients were averaged to estimate the mean rates and associated standard deviation.

### Coarse-grained Langevin dynamics simulations

The CG Langevin dynamics simulations of the DRYG-CTR and CTR constructs were carried out using OpenMM version 7.5^78^. Each amino acid was represented as a single bead mapped to the Cα carbon of each residue. Molecular interactions between beads were accounted for as bonded or non-bonded. Bonded interactions were considered for two beads directly linked by a harmonic potential, with an equilibrium bond length of 0.38 nm and a force constant of 8033 kJ mol^-1^ nm^-2^. Non-bonded interactions were modeled for all beads not directly bonded using the interaction potentials implemented in CALVADOS2^79,80^. Particularly, dispersion forces were accounted for by a truncated and shifted Ashbaugh-Hatch potential, in which a residue-specific hydrophobicity parameter λ, scales interaction strengths between beads. Electrostatic forces acting between charged particles were accounted for, as in implicit solvent, by a salt-screening Debye-Hückel based potential, truncated at a cut-off distance of 2 nm. In this model, beads representing aspartate and glutamate carry a -1 charge, while lysine and arginine beads carry a +1 charge.

Single chains of DRYG-CTR and CTR were each modeled as linear chains and placed in a box with dimensions 100 nm^3^. To simulate the oligomeric systems, we followed the slab method suggested to well simulate IDPs in the condensed phase^81^. 50 chains of the DRYG-CTR construct were modeled as linear chains, equally spaced and placed in a triclinic box measuring 25 x 25 x 200 nm along x, y and z, respectively. All systems were simulated considering periodic boundary conditions and at two ionic strengths: 0.117 and 0.192 M. After energy minimization, each system was simulated at a temperature of 293 K, with a time step of 0.01 ps and a friction coefficient of 0.01 ps^-1^, saving coordinates every 1 ns. Each system was simulated for a total of 10 µs with the first μs discarded as considered equilibration time.

Using the Förster equation (Eq. 4), transfer efficiencies were calculated from the simulated distances between the experimentally labeled residues along the protein, implementing a Förster radius of 5.9 nm, reflective of the *R*_0_ of the Cy3B/CF660R dye-pair used in the smFRET experiments. All the simulations were performed without explicitly modeling FRET dyes, as those were not parameterized within the used coarse-grained model. Nevertheless, the effect of dyes on the retrieved ensembles has been shown in multiple studies^45,46,48^ to be negligible. The agreement between experimental and computed FRET efficiencies was obtained by calculating the concordance correlation coefficient of the two datasets as follows:

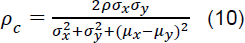

where *μ_x_* and *μ_y_* are the means of the experimental and computational FRET efficiencies, respectively, while *σ_x_* and *σ_y_* are the standard deviations of each dataset.

The collected trajectories were analyzed using a combination of *in-house* scripts, MDAnalysis libraries^82^ and tools available in GROMACS version 2021.6^83^. Clustering was performed using gmx clustsize, with a distance cutoff of 2 nm. For each simulation step, two or more chains were considered bound if the distance between two or more beads fell within 1 nm. Intramolecular and intermolecular contacts were calculated from the single-chain and oligomeric systems, respectively. Intermolecular contacts were obtained by firstly calculating the average distance between all beads of a bound cluster: distance arrays were then binarized to create a Boolean contact matrix (1 if bound, 0 if not bound) considering a cutoff of 1 nm and the final fraction of residue-wide contacts was calculated as a time-average over the simulated trajectory. The map of the fraction of contacts gained for oligomers (shown in Fig. 6f) was quantified by subtracting the intra-chain contacts sampled in the single chain simulations from the inter-chain contacts sampled in the oligomeric state. Representative snapshots of the simulated ensembles were obtained using VMD version 1.9.3^84^.

## Data availability

All data supporting the findings of this work are available within the main text and its Supplementary Information. The gene sequence of eIF4B used in this work is provided in Supplementary Table 1. The list of oligonucleotide primers used in this work is provided in Supplementary Table 2. The list of sequences for protein constructs used in this work is provided in Supplementary Table 3.

The raw data supporting the findings of this study are available from the corresponding authors upon request. The chemical shift assignments for eIF4B-CTR were previously deposited in the BMRB (https://bmrb.io/) under accession codes 51952, 51953 and 51957.

The CG simulation trajectories are available from: https://doi.org/10.5281/zenodo.12204791

## Acknowledgments

We thank Benjamin Schuler and Daniel Nettels for support of the smFRET data analysis package Fretica. We thank Jean-Michel Blanc and the IECB biochemistry facility for assisting the preparation of the wild-type eIF4B plasmid. We thank Axelle Grélard and Estelle Morvan for assistance and access to the NMR spectrometers in the structural biophysico-chemistry facility at the European Institute of Chemistry and Biology (IECB; CNRS UAR 3033, Inserm US001). D.M. thanks the faculty of Science of the University of Auckland for providing the computational facilities used for the simulations presented in this work.

This work was supported by grants from the Fondation pour la Recherche Médicale (AJE20171239128 to M.A.), Junior Chair IdEx Univ. Bordeaux (OPE-2018-0405 to M.A.), ANR JCJC (DISTINCT: ANR-20-CE11-0007 to M.A.), Region of Nouvelle-Aquitaine (grant AAPR2020-2020-8589310 to M.A.) and INSERM (M.A., C.D.M). B.C.S was supported by the FRM postdoctoral fellowship (SPF202110014130). V.U. was supported by a PhD scholarship of the Faculty of Science, The University of Auckland.

## Author information

V. Esperance Aho

Present address: Institut de Biologie Structurale (IBS), UMR 5075, F-38044 Grenoble, France These authors contributed equally: Bikash Chandra Swain, Pascale Sarkis and Vanessa Ung.

## Authors and affiliations

Univ. Bordeaux, Inserm U1212, CNRS UMR 5320, ARNA, Institut Européen de Chimie et Biologie, F-33600 Pessac, France

Bikash Chandra Swain, Pascale Sarkis, Sabrina Rousseau, Laurent Fernandez, Ani Meltonyan, V. Esperance Aho, Cameron D. Mackereth, Mikayel Aznauryan

School of Chemical Science, University of Auckland, Auckland, New Zealand Vanessa Ung, Davide Mercadante

## Contributions

M.A. and C.D.M. conceived and designed the experiments; B.C.S., P.S., S.R., L.F. A.M. V.E.A, C.D.M and M.A. performed the experiments; D.M. conceived and designed the simulations; V.U. performed the simulations; B.C.S., P.S., V.U., L.F., D.M., C.D.M and M.A. analyzed the data; M.A. wrote the paper with the contributions from B.C.S., P.S., V.U., S.R., D.M. and C.D.M.

## Corresponding authors

Correspondence to: Mikayel Aznauryan, Cameron D. Mackereth

## References

1 Jackson, R. J., Hellen, C. U. & Pestova, T. V. The mechanism of eukaryotic translation initiation and principles of its regulation. Nat Rev Mol Cell Biol 11, 113–127 (2010).

2 Sokabe, M. & Fraser, C. S. Toward a Kinetic Understanding of Eukaryotic Translation. Cold Spring Harb Perspect Biol 11 (2019).

3 Marintchev, A. et al. Topology and regulation of the human eIF4A/4G/4H helicase complex in translation initiation. Cell 136, 447–460 (2009).

4 Hinnebusch, A. G. The scanning mechanism of eukaryotic translation initiation. Annu Rev Biochem 83, 779–812 (2014).

5 Merrick, W. C. eIF4F: a retrospective. J Biol Chem 290, 24091–24099 (2015).

6 Brito Querido, J., et al. The structure of a human translation initiation complex reveals two independent roles for the helicase eIF4A. Nat Struct Mol Biol 31, 455–464 (2024).

7 Ozes, A. R., Feoktistova, K., Avanzino, B. C. & Fraser, C. S. Duplex unwinding and ATPase activities of the DEAD-box helicase eIF4A are coupled by eIF4G and eIF4B. J Mol Biol 412, 674–687 (2011).

8 Nielsen, K. H. et al. Synergistic activation of eIF4A by eIF4B and eIF4G. Nucleic Acids Res 39, 2678–2689 (2011).

9 Rozovsky, N., Butterworth, A. C. & Moore, M. J. Interactions between eIF4AI and its accessory factors eIF4B and eIF4H. RNA 14, 2136–2148 (2008).

10 Garcia-Garcia, C., Frieda, K. L., Feoktistova, K., Fraser, C. S. & Block, S. M. RNA BIOCHEMISTRY. Factor-dependent processivity in human eIF4A DEAD-box helicase. Science 348, 1486–1488 (2015).

11 Methot, N., Pause, A., Hershey, J. W. & Sonenberg, N. The translation initiation factor eIF-4B contains an RNA-binding region that is distinct and independent from its ribonucleoprotein consensus sequence. Mol Cell Biol 14, 2307–2316 (1994).

12 Rogers, G. W., Jr., Richter, N. J., Lima, W. F. & Merrick, W. C. Modulation of the helicase activity of eIF4A by eIF4B, eIF4H, and eIF4F. J Biol Chem 276, 30914–30922 (2001).

13 Aitken, C. E. & Lorsch, J. R. A mechanistic overview of translation initiation in eukaryotes. Nat Struct Mol Biol 19, 568–576 (2012).

14 Brito Querido, J., et al. Structure of a human 48S translational initiation complex. Science 369, 1220–1227 (2020).

15 Brito Querido, J., Diaz-Lopez, I. & Ramakrishnan, V. The molecular basis of translation initiation and its regulation in eukaryotes. Nat Rev Mol Cell Biol 25, 168–186 (2024).

16 Hashem, Y. & Frank, J. The Jigsaw Puzzle of mRNA Translation Initiation in Eukaryotes: A Decade of Structures Unraveling the Mechanics of the Process. Annu Rev Biophys 47, 125–151 (2018).

17 Simonetti, A., Guca, E., Bochler, A., Kuhn, L. & Hashem, Y. Structural Insights into the Mammalian Late-Stage Initiation Complexes. Cell Rep 31, 107497 (2020).

18 des Georges, A. et al. Structure of mammalian eIF3 in the context of the 43S preinitiation complex. Nature 525, 491–495 (2015).

19 Eliseev, B. et al. Structure of a human cap-dependent 48S translation pre-initiation complex. Nucleic Acids Res 46, 2678–2689 (2018).

20 Shahbazian, D. et al. Control of cell survival and proliferation by mammalian eukaryotic initiation factor 4B. Mol Cell Biol 30, 1478–1485 (2010).

21 Wang, Y. et al. Mitotic MELK-eIF4B signaling controls protein synthesis and tumor cell survival. Proc Natl Acad Sci U S A 113, 9810–9815 (2016).

22 Shahbazian, D., Parsyan, A., Petroulakis, E., Hershey, J. & Sonenberg, N. eIF4B controls survival and proliferation and is regulated by proto-oncogenic signaling pathways. Cell Cycle 9, 4106–4109 (2010).

23 Sen, N. D., Zhou, F., Harris, M. S., Ingolia, N. T. & Hinnebusch, A. G. eIF4B stimulates translation of long mRNAs with structured 5’ UTRs and low closed-loop potential but weak dependence on eIF4G. Proc Natl Acad Sci U S A 113, 10464–10472 (2016).

24 Horvilleur, E. et al. A role for eukaryotic initiation factor 4B overexpression in the pathogenesis of diffuse large B-cell lymphoma. Leukemia 28, 1092–1102 (2014).

25 Buchan, J. R. & Parker, R. Eukaryotic stress granules: the ins and outs of translation. Mol Cell 36, 932–941 (2009).

26 Low, W. K. et al. Inhibition of eukaryotic translation initiation by the marine natural product pateamine A. Mol Cell 20, 709–722 (2005).

27 Mokas, S. et al. Uncoupling stress granule assembly and translation initiation inhibition. Mol Biol Cell 20, 2673–2683 (2009).

28 Fleming, K. et al. Solution structure and RNA interactions of the RNA recognition motif from eukaryotic translation initiation factor 4B. Biochemistry 42, 8966–8975 (2003).

29 Mondal, S. et al. Backbone resonance assignments of the C-terminal region of human translation initiation factor eIF4B. Biomol NMR Assign 17, 199–203 (2023).

30 Naranda, T., Strong, W. B., Menaya, J., Fabbri, B. J. & Hershey, J. W. Two structural domains of initiation factor eIF-4B are involved in binding to RNA. J Biol Chem 269, 14465–14472 (1994).

31 Milburn, S. C., Hershey, J. W., Davies, M. V., Kelleher, K. & Kaufman, R. J. Cloning and expression of eukaryotic initiation factor 4B cDNA: sequence determination identifies a common RNA recognition motif. EMBO J 9, 2783–2790 (1990).

32 Methot, N., Song, M. S. & Sonenberg, N. A region rich in aspartic acid, arginine, tyrosine, and glycine (DRYG) mediates eukaryotic initiation factor 4B (eIF4B) self-association and interaction with eIF3. Mol Cell Biol 16, 5328–5334 (1996).

33 Raught, B. et al. Phosphorylation of eucaryotic translation initiation factor 4B Ser422 is modulated by S6 kinases. EMBO J 23, 1761–1769 (2004).

34 Erdos, G., Pajkos, M. & Dosztanyi, Z. IUPred3: prediction of protein disorder enhanced with unambiguous experimental annotation and visualization of evolutionary conservation. Nucleic Acids Res 49, W297–W303 (2021).

35 O’Brien, K. T., Mooney, C., Lopez, C., Pollastri, G. & Shields, D. C. Prediction of polyproline II secondary structure propensity in proteins. R Soc Open Sci 7, 191239 (2020).

36 Hoffmann, A. et al. Mapping protein collapse with single-molecule fluorescence and kinetic synchrotron radiation circular dichroism spectroscopy. Proc Natl Acad Sci U S A 104, 105–110 (2007).

37 Aznauryan, M. et al. Comprehensive structural and dynamical view of an unfolded protein from the combination of single-molecule FRET, NMR, and SAXS. Proc Natl Acad Sci U S A 113, E5389–5398 (2016).

38 Schuler, B., Soranno, A., Hofmann, H. & Nettels, D. Single-Molecule FRET Spectroscopy and the Polymer Physics of Unfolded and Intrinsically Disordered Proteins. Annu Rev Biophys 45, 207–231 (2016).

39 Schuler, B. Perspective: Chain dynamics of unfolded and intrinsically disordered proteins from nanosecond fluorescence correlation spectroscopy combined with single-molecule FRET. J Chem Phys 149, 010901 (2018).

40 Martin, E. W. et al. Sequence Determinants of the Conformational Properties of an Intrinsically Disordered Protein Prior to and upon Multisite Phosphorylation. J Am Chem Soc 138, 15323–15335 (2016).

41 Reiersen, H. & Rees, A. R. The hunchback and its neighbours: proline as an environmental modulator. Trends Biochem Sci 26, 679–684 (2001).

42 Soranno, A. et al. Quantifying internal friction in unfolded and intrinsically disordered proteins with single-molecule spectroscopy. Proc Natl Acad Sci U S A 109, 17800–17806 (2012).

43 Record, M. T., Jr., Anderson, C. F. & Lohman, T. M. Thermodynamic analysis of ion effects on the binding and conformational equilibria of proteins and nucleic acids: the roles of ion association or release, screening, and ion effects on water activity. Q Rev Biophys 11, 103–178 (1978).

44 Hoffmann, A. et al. Quantifying heterogeneity and conformational dynamics from single molecule FRET of diffusing molecules: recurrence analysis of single particles (RASP). Phys Chem Chem Phys 13, 1857–1871 (2011).

45 Borgia, A. et al. Extreme disorder in an ultrahigh-affinity protein complex. Nature 555, 61–66 (2018).

46 Bjarnason, S. et al. DNA binding redistributes activation domain ensemble and accessibility in pioneer factor Sox2. Nat Commun 15, 1445 (2024).

47 Holmstrom, E. D., Liu, Z., Nettels, D., Best, R. B. & Schuler, B. Disordered RNA chaperones can enhance nucleic acid folding via local charge screening. Nat Commun 10, 2453 (2019).

48 Heidarsson, P. O. et al. Release of linker histone from the nucleosome driven by polyelectrolyte competition with a disordered protein. Nat Chem 14, 224–231 (2022).

49 Guillen-Boixet, J. et al. RNA-Induced Conformational Switching and Clustering of G3BP Drive Stress Granule Assembly by Condensation. Cell 181, 346–361 e317 (2020).

50 Bremer, A. et al. Deciphering how naturally occurring sequence features impact the phase behaviours of disordered prion-like domains. Nat Chem 14, 196–207 (2022).

51 Dao, T. P., Rajendran, A., Galagedera, S. K. K., Haws, W. & Castaneda, C. A. Short disordered termini and proline-rich domain are major regulators of UBQLN1/2/4 phase separation. Biophys J (2023).

52 Itzhak, D. N., Tyanova, S., Cox, J. & Borner, G. H. Global, quantitative and dynamic mapping of protein subcellular localization. Elife 5 (2016).

53 Hein, M. Y. et al. A human interactome in three quantitative dimensions organized by stoichiometries and abundances. Cell 163, 712–723 (2015).

54 Duncan, R. & Hershey, J. W. Identification and quantitation of levels of protein synthesis initiation factors in crude HeLa cell lysates by two-dimensional polyacrylamide gel electrophoresis. J Biol Chem 258, 7228–7235 (1983).

55 Altmann, M., Wittmer, B., Methot, N., Sonenberg, N. & Trachsel, H. The Saccharomyces cerevisiae translation initiation factor Tif3 and its mammalian homologue, eIF-4B, have RNA annealing activity. EMBO J 14, 3820–3827 (1995).

56 Marzahn, M. R. et al. Higher-order oligomerization promotes localization of SPOP to liquid nuclear speckles. EMBO J 35, 1254–1275 (2016).

57 Dao, T. P. et al. Ubiquitin Modulates Liquid-Liquid Phase Separation of UBQLN2 via Disruption of Multivalent Interactions. Mol Cell 69, 965–978 e966 (2018).

58 Kar, M. et al. Phase-separating RNA-binding proteins form heterogeneous distributions of clusters in subsaturated solutions. Proc Natl Acad Sci U S A 119, e2202222119 (2022).

59 Zhao, H. et al. Energetic and structural features of SARS-CoV-2 N-protein co-assemblies with nucleic acids. iScience 24, 102523 (2021).

60 Ray, S. et al. Mass photometric detection and quantification of nanoscale alpha-synuclein phase separation. Nat Chem 15, 1306–1316 (2023).

61 Nosella, M. L. & Forman-Kay, J. D. Phosphorylation-dependent regulation of messenger RNA transcription, processing and translation within biomolecular condensates. Curr Opin Cell Biol 69, 30–40 (2021).

62 Hofweber, M. & Dormann, D. Friend or foe-Post-translational modifications as regulators of phase separation and RNP granule dynamics. J Biol Chem 294, 7137–7150 (2019).

63 Snead, W. T. & Gladfelter, A. S. The Control Centers of Biomolecular Phase Separation: How Membrane Surfaces, PTMs, and Active Processes Regulate Condensation. Mol Cell 76, 295–305 (2019).

64 Watanabe, K. et al. Cells recognize osmotic stress through liquid-liquid phase separation lubricated with poly(ADP-ribose). Nat Commun 12, 1353 (2021).

65 Gao, C. et al. Hyperosmotic-stress-induced liquid-liquid phase separation of ALS-related proteins in the nucleus. Cell Rep 40, 111086 (2022).

66 Franzmann, T. M. & Alberti, S. Protein Phase Separation as a Stress Survival Strategy. Cold Spring Harb Perspect Biol 11 (2019).

67 Jalihal, A. P. et al. Multivalent Proteins Rapidly and Reversibly Phase-Separate upon Osmotic Cell Volume Change. Mol Cell 79, 978–990 e975 (2020).

68 Konig, I. et al. Single-molecule spectroscopy of protein conformational dynamics in live eukaryotic cells. Nat Methods 12, 773–779 (2015).

69 Milkovic, N. M. & Mittag, T. Determination of Protein Phase Diagrams by Centrifugation. Methods Mol Biol 2141, 685–702 (2020).

70 Delaglio, F. et al. NMRPipe: a multidimensional spectral processing system based on UNIX pipes. J Biomol NMR 6, 277–293 (1995).

71 Muller, B. K., Zaychikov, E., Brauchle, C. & Lamb, D. C. Pulsed interleaved excitation. Biophys J 89, 3508–3522 (2005).

72 Cubuk, J. et al. The SARS-CoV-2 nucleocapsid protein is dynamic, disordered, and phase separates with RNA. Nat Commun 12, 1936 (2021).

73 Schuler, B., Muller-Spath, S., Soranno, A. & Nettels, D. Application of confocal single-molecule FRET to intrinsically disordered proteins. Methods Mol Biol 896, 21–45 (2012).

74 Holmstrom, E. D. et al. Accurate Transfer Efficiencies, Distance Distributions, and Ensembles of Unfolded and Intrinsically Disordered Proteins From Single-Molecule FRET. Methods Enzymol 611, 287–325 (2018).

75 Zheng, W. et al. Inferring properties of disordered chains from FRET transfer efficiencies. J Chem Phys 148, 123329 (2018).

76 Nettels, D., Hoffmann, A. & Schuler, B. Unfolded protein and peptide dynamics investigated with single-molecule FRET and correlation spectroscopy from picoseconds to seconds. J Phys Chem B 112, 6137–6146 (2008).

77 Gopich, I. V., Nettels, D., Schuler, B. & Szabo, A. Protein dynamics from single-molecule fluorescence intensity correlation functions. J Chem Phys 131, 095102 (2009).

78 Eastman, P. et al. OpenMM 7: Rapid development of high performance algorithms for molecular dynamics. PLoS Comput Biol 13, e1005659 (2017).

79 Tesei, G., Schulze, T. K., Crehuet, R. & Lindorff-Larsen, K. Accurate model of liquid-liquid phase behavior of intrinsically disordered proteins from optimization of single-chain properties. Proc Natl Acad Sci U S A 118, e2111696118 (2021).

80 Tesei, G. & Lindorff-Larsen, K. Improved predictions of phase behaviour of intrinsically disordered proteins by tuning the interaction range [version 2; peer review: 2 approved]. Open Research Europe 2 (2023).

81 Dignon, G. L., Zheng, W., Best, R. B., Kim, Y. C. & Mittal, J. Relation between single-molecule properties and phase behavior of intrinsically disordered proteins. Proc Natl Acad Sci U S A 115, 9929–9934 (2018).

82 Michaud-Agrawal, N., Denning, E. J., Woolf, T. B. & Beckstein, O. MDAnalysis: a toolkit for the analysis of molecular dynamics simulations. J Comput Chem 32, 2319–2327 (2011).

83 GROMACS 2021 Source code.

84 Humphrey, W., Dalke, A. & Schulten, K. VMD: visual molecular dynamics. J Mol Graph 14, 33–38, 27-38 (1996).

