## Supplementary Information for "Disordered regions of human eIF4B orchestrate a dynamic self-association landscape"

### Disordered region of human eIF4B orchestrates a dynamic balance across self-association landscape

[#] these authors contributed equally

**Corresponding authors:*

Cameron D. Mackereth

ORCID : 0000-0002-0776-7947

Mikayel Aznauryan

ORCID : 0000-0002-5395-3441

### Additional methods

#### Purification of the DRYG construct

Bacteria were harvested by centrifugation and then resuspended in a lysis buffer consisting of 50 mM Tris-HCl (pH 7.8), 250 mM KCl, 100 mg/L lysozyme, 1 mM PMSF, 2 mM β-mercaptoethanol, and one protease inhibitor cocktail tablet (Roche). Following the sonication on ice (1 min at 40% power, alternating 20s of sonication and 20s of pause), the sample was centrifuged at 20000 x *g* for 40 min at 4 °C. The pellet was resuspended in washing buffer consisting of 50 mM Tris-HCl (pH 7.8), 250 mM KCl, 1% Triton-X100, 2 mM β-mercaptoethanol, 1 M urea. After another round of sonication on ice (1 min at 40% power, alternating 20s of sonication and 20s of pause), the sample was again centrifuged at 20000 x *g* for 20 min at 4 °C. This step was repeated until the supernatant was cleared. To remove the Triton-X100, a final washing was performed with the washing buffer without Triton-X100. After centrifugation, the protein was extracted by solubilizing the pellet in the GdmCl-containing buffer consisting of 50 mM Tris-HCl (pH 7.8), 8 M GdmCl, 250 mM KCl, 30mM imidazole and 2 mM β-mercaptoethanol. This step was followed by an additional round of sonication on ice (1 min at 40% power, alternating 20s of sonication and 20s of pause). This mixture was left for one hour at room temperature with stirring, and was thereafter filtered with a 0.7 µm glass Microfilter GF/F (GE Healthcare Life Sciences Whatman). This protein was largely pure and devoid of any shorter-length protein degradation fragments. The His-tag of this construct was not cut due to inefficient cleavage reaction. The protein was then purified with reverse-phase HPLC on a C18 column (ReproSil Gold 200, Dr. Maisch), followed by lyophilization.

**Supplementary table 1: The codon-optimized eIF4B gene sequence used in this work**

```
          atgaaatcttctcaccatcaccatcaccatgaaaacctgtacttccaagga
           M  K  S  S  H  H  H  H  H  H  E  N  L  Y  F  Q  G His-tag
atggcggcgagcgcgaaaaagaaaaacaaaaaaggcaaaaccattagcctgaccgatttt
 M  A  A  S  A  K  K  K  N  K  K  G  K  T  I  S  L  T  D  F  20
ctggcggaagatggcggcaccggcggcggcagcacctatgtgagcaaaccggtgagctgg
 L  A  E  D  G  G  T  G  G  G  S  T  Y  V  S  K  P  V  S  W 40
gcggatgaaaccgatgatctggaaggcgatgtgagcaccacctggcatagcaacgatgat
 A  D  E  T  D  D  L  E  G  D  V  S  T  T  W  H  S  N  D  D  60
gatgtgtatcgcgcgccgccgattgatcgcagcattctgccgaccgcgccgcgcgcggcg
 D  V  Y  R  A  P  P  I  D  R  S  I  L  P  T  A  P  R  A  A  80
cgcgaaccgaacattgatcgcagccgcctgccgaaaagcccgccgtataccgcgtttctg
 R  E  P  N  I  D  R  S  R  L  P  K  S  P  P  Y  T  A  F  L  100
ggcaacctgccgtatgatgtgaccgaagaaagcattaaagaatttttcgcggcctgaac
 G  N  L  P  Y  D  V  T  E  E  S  I  K  E  F  F  R  G  L  N  RRM domain
attagcgcggtgcgcctgccgcgcgaaccgagcaacccggaacgcctgaaaggctttggc
 I  S  A  V  R  L  P  R  E  P  S  N  P  E  R  L  K  G  F  G  140
tatgcggaatttgaagatctggatagcctgctgagcgcgctgagcctgaacgaagaaagc
 Y  A  E  F  E  D  L  D  S  L  L  S  A  L  S  L  N  E  E  S  160
ctgggcaaccgccgcattcgcgtggatgtggcggatcaggcgcaggataaagatcgcgat
 L  G  N  R  R  I  R  V  D  V  A  D  Q  A  Q  D  K  D  R  D  180
gatcgcagctttggccgcgatcgcaaccgcgatagcgataaaaccgataccgattggcgc
 D  R  S  F  G  R  D  R  N  R  D  S  D  K  T  D  T  D  W  R  200
gcgcgcccggcgaccgatagctttgatgattatccgccgcgccgcggcgatgatagcttt
 A  R  P  A  T  D  S  F  D  D  Y  P  P  R  R  G  D  D  S  F  220
ggcgataaatatcgcgatcgctatgatagcgatcgctatcgcgatggctatcgcgatggc
 G  D  K  Y  R  D  R  Y  D  S  D  R  Y  R  D  G  Y  R  D  G  240
tatcgcgatggcccgcgccgcgatatggatcgctatggcggccgcgatcgctatgatgat
 Y  R  D  G  P  R  R  D  M  D  R  Y  G  G  R  D  R  Y  D  D  DRYG
cgcggcagccgcgattatgatcgcggctatgatagccgcattggcagcggccgccgcgcg
 R  G  S  R  D  Y  D  R  G  Y  D  S  R  I  G  S  G  R  R  A  280
tttggcagcggctatcgccgcgatgatgattatcgcggcggcggcgatcgctatgaagat
 F  G  S  G  Y  R  R  D  D  D  Y  R  G  G  G  D  R  Y  E  D  300
cgctatgatcgccgcgatgatcgcagctggagcagccgcgatgattatagccgcgatgat
 R  Y  D  R  R  D  D  R  S  W  S  S  R  D  D  Y  S  R  D  D  320
tatcgccgcgatgatcgcggcccgccgcagcgcccgaaactgaacctgaaaccgcgcagc
 Y  R  R  D  D  R  G  P  P  Q  R  P  K  L  N  L  K  P  R  S  330
accccgaaagaagatgatagcagcgcgagcaccagccagagcacccgcgcggcgagcatt
 T  P  K  E  D  D  S  S  A  S  T  S  Q  S  T  R  A  A  S  I  360
tttggcggcgcgaaaccggtggataccgcggcgcgcgaacgcgaagtggaagaacgcctg
 F  G  G  A  K  P  V  D  T  A  A  R  E  R  E  V  E  E  R  L  ARM
cagaaagaacaggaaaaactgcagcgccagctggatgaaccgaaactggaacgccgcccg
 Q  K  E  Q  E  K  L  Q  R  Q  L  D  E  P  K  L  E  R  R  P  400
cgcgaacgccatccgagctggcgcagcgaagaaacccaggaacgcgaacgcagccgcacc
 R  E  R  H  P  S  W  R  S  E  E  T  Q  E  R  E  R  S  R  T  420
ggcagcgaaagcagccagaccggcaccagcaccaccagcagccgcaacgcgcgccgccgc
 G  S  E  S  S  Q  T  G  T  S  T  T  S  S  R  N  A  R  R  R  440
gaaagcgaaaaaagcctggaaaacgaaaccctgaacaaagaagaagattgccatagcccg
 E  S  E  K  S  L  E  N  E  T  L  N  K  E  E  D  C  H  S  P  457
accagcaaaccgccgaaaccggatcagccgctgaaagtgatgccggcgccgccgccgaaa
 T  S  K  P  P  K  P  D  Q  P  L  K  V  M  P  A  P  P  P  K  480
gaaaacgcgtgggtgaaacgcagcagcaacccgccggcgcgcagccagagcagcgatacc
 E  N  A  W  V  K  R  S  S  N  P  P  A  R  S  Q  S  S  D  T  500
gaacagcagagcccgaccagcggcggcggcaaagtggcgccggcgcagccgagcgaagaa
 E  Q  Q  S  P  T  S  G  G  G  K  V  A  P  A  Q  P  S  E  E  520
ggcccgggccgcaaagatgaaaacaaagtggatggcatgaacgcgccgaaaggccagacc
 G  P  G  R  K  D  E  N  K  V  D  G  M  N  A  P  K  G  Q  T  540
ggcaacagcagccgcggcccgggcgatggcggcaaccgcgatcattggaaagaaagcgat
```

G N S S R G P G D G G N R D H W K E S D 560
cgcaaagatggcaaaaaagatcaggatagccgcagcgcgccggaaccgaaaaaaaccggaa
R K D G K K D Q D S R S A P E P K K P E 580
gaaaacccggcgagcaaatttagcagcgcgagcaaatatgcggcgctgagcgtggatggc
E N P A S K F S S A S K Y A A L S V D G 600
gaagatgaaaacgaaggcgaagattatgcggaataataacattggaagtggataacggat
E D E N E G E D Y A E

**Supplementary table 2: The list of primers used in this work**

Primers to create truncated constructs:

| Name | Sequence | truncated construct |
|---|---|---|
| eIF4B-Δ458-611-F | GAAGATTGCTAACATTGGAAGTGG | CTR-N;<br>DRYG-CTR-N |
| eIF4B-Δ458-611-R | CTTCCAATGTTAGCAATCTTCTTCTTTG | |
| eIF4B-C-Δ-332-457-F | CTTCCAAGGCCATAGCCCGACCAGCAAA | CTR-C |
| eIF4B-C-Δ-332-457-R | CGGGCTATGGCCTTGGAAGTACAGGTTTTC | |
| eIF4B-Δ-332-457-F | GCCCACCTCAAAGACATAGCCCGACCAGCAAACCGC | DRYG-CTR-C |
| eIF4B-Δ-332-457-R | TCTTTGAGGTGGGCCGCGATCATCGCGGCG | |
| eIF4B- DRYG-C332-F | GCCGCAGCGCTGCTAATAACATTGGAAGTGGATAACGG | DRYG |
| eIF4B-DRYG-C332-R | AATGTTATTAGCAGCGCTGCGGCGGGCC | |

Primers for cysteine mutants:

| Name | Sequence | applicable to |
|---|---|---|
| eIF4B-F361C-F | CGAGCATTTGTGGCGGCGCGAAAC | CTR;<br>DRYG-CTR |
| eIF4B-F361C-R | TCGCGCCGCCACAAATGCTCGC | |
| eIF4B-G523C-F | GAAGAAGGCCCGTGCCGCAAAGATGAAAAC | CTR;<br>DRYG-CTR |
| eIF4B-G523C-R | CATCTTTGCGGCACGGGCCTTCTTCGCTCGG | |
| eIF4B-C457A-F | AGAAGAAGATGCCCATAGCCCGACCAGCAAAC | |

| | | |
|---|---|---|
| eIF4B-C457A-R | GGTCGGGCTATGGGCATCTTCTTCTTTGTTCAGGG | CTR; DRYG-CTR |
| eIF4B-Y609C-F | GAAGGCGAAGATTGTGCGGAATAATAACATTGG | CTR; DRYG-CTR |
| eIF4B-Y609C-R | GTTATTATTCCGCACAATCTTCGCCTTCGTTTTCATCTTC | |
| eIF4B-W407C-F | CCGAGCTGTCGCAGCGAAGAAAC | CTR; DRYG-CTR |
| eIF4B-W407C-R | TCTTCGCTGCGACAGCTCGGATG | |
| eIF4B-C332P-F | CGCAGCGCCCCAAACTGAACCTGAAA | DRYG-CTR |
| eIF4B-C332P-R | TTCAGTTTGGGGCGCTGCGGC | |
| eIF4B-DRYG-P213C-R | GCTGTCGCCGCGGCGAT | DRYG-CTR |
| eIF4B-DRYG-P213C-F | CCGCGGCGACAGCCTTGGAAGTACAGGTTT | |
| eIF4B-CTR-C332P-F | CTTCCAAGGCCCCAAACTGAACCTGAAACC | CTR |
| eIF4B-CTR-C332P-R | GGTTCAGTTTGGGGCCTTGGAAGTACAGG | |

### Supplementary table 2: The sequences of the eIF4B constructs used in this work

| **DRYG-CTR constructs** | | |
|---|---|---|
| Construct | Labeling sites | Sequence |
| **DRYG-CTR**$_{\mathbf{P213C-P332C}}$ | **213-332** | 220 230 240 250 260 270<br>GCRRGDDSF GDKYRDRYDS DRYRDGYRDG YRDGPRRDMD RYGGRDRYDD RGSRDYDRGY<br>280 290 300 310 320 330<br>DSRIGSGRRA FGSGYRRDDD YRGGGDRYED RYDRRDDRSW SSRDDYSRDD YRRDDRGPPQ<br>340 350 360 370 380 390<br>RCKLNLKPRS TPKEDDSSAS TSQSTRAASI FGGAKPVDTA AREREVEERL QKEQEKLQRQ<br>400 410 420 430 440 450<br>LDEPKLERRP RERHPSWRSE ETQERERSRT GSESSQTGTS TTSSRNARRR ESEKSLENET<br>460 470 480 490 500 510<br>LNKEED**A**HSP TSKPPKPDQP LKVMPAPPPK ENAWVKRSSN PPARSQSSDT EQQSPTSGGG<br>520 530 540 550 560 570<br>KVAPAQPSEE GPGRKDENKV DGMNAPKGQT GNSSRGPGDG GNRDHWKESD RKDGKKDQDS<br>580 590 600 610<br>RSAPEPKKPE ENPASKFSSA SKYAALSVDG EDENEGEDYA E |
| **DRYG-CTR**$_{\mathbf{P332C-C457}}$ [1] | **332-457** | 220 230 240 250 260 270<br>GRRGDDSF GDKYRDRYDS DRYRDGYRDG YRDGPRRDMD RYGGRDRYDD RGSRDYDRGY<br>280 290 300 310 320 330<br>DSRIGSGRRA FGSGYRRDDD YRGGGDRYED RYDRRDDRSW SSRDDYSRDD YRRDDRGPPQ<br>340 350 360 370 380 390<br>RCKLNLKPRS TPKEDDSSAS TSQSTRAASI FGGAKPVDTA AREREVEERL QKEQEKLQRQ<br>400 410 420 430 440 450<br>LDEPKLERRP RERHPSWRSE ETQERERSRT GSESSQTGTS TTSSRNARRR ESEKSLENET<br>460 470 480 490 500 510<br>LNKEEDCHSP TSKPPKPDQP LKVMPAPPPK ENAWVKRSSN PPARSQSSDT EQQSPTSGGG |

[1] This DRYG-CTR construct is used for NMR and CD experiments

| | | |
|---|---|---|
| | | 520 530 540 550 560 570<br>KVAPAQPSEE GPGRKDENKV DGMNAPKGQT GNSSRGPGDG GNRDHWKESD RKDGKKDQDS<br>580 590 600 610<br>RSAPEPKKPE ENPASKFSSA SKYAALSVDG EDENEGEDYA E |
| **DRYG-CTR$_{C457\text{-}G523C}$** | **457-523** | 220 230 240 250 260 270<br>GRRGDDSF GDKYRDRYDS DRYRDGYRDG YRDGPRRDMD RYGGRDRYDD RGSRDYDRGY<br>280 290 300 310 320 330<br>DSRIGSGRRA FGSGYRRDDD YRGGGDRYED RYDRRDDRSW SSRDDYSRDD YRRDDRGPPQ<br>340 350 360 370 380 390<br>RPKLNLKPRS TPKEDDSSAS TSQSTRAASI FGGAKPVDTA AREREVEERL QKEQEKLQRQ<br>400 410 420 430 440 450<br>LDEPKLERRP RERHPSWRSE ETQERERSRT GSESSQTGTS TTSSRNARRR ESEKSLENET<br>460 470 480 490 500 510<br>LNKEED**C**HSP TSKPPKPDQP LKVMPAPPPK ENAWVKRSSN PPARSQSSDT EQQSPTSGGG<br>520 530 540 550 560 570<br>KVAPAQPSEE GP**C**RKDENKV DGMNAPKGQT GNSSRGPGDG GNRDHWKESD RKDGKKDQDS<br>580 590 600 610<br>RSAPEPKKPE ENPASKFSSA SKYAALSVDG EDENEGEDYA E |
| **DRYG-CTR$_{G523C\text{-}Y609C}$** | **523-609** | 220 230 240 250 260 270<br>GRRGDDSF GDKYRDRYDS DRYRDGYRDG YRDGPRRDMD RYGGRDRYDD RGSRDYDRGY<br>280 290 300 310 320 330<br>DSRIGSGRRA FGSGYRRDDD YRGGGDRYED RYDRRDDRSW SSRDDYSRDD YRRDDRGPPQ<br>340 350 360 370 380 390<br>RPKLNLKPRS TPKEDDSSAS TSQSTRAASI FGGAKPVDTA AREREVEERL QKEQEKLQRQ<br>400 410 420 430 440 450<br>LDEPKLERRP RERHPSWRSE ETQERERSRT GSESSQTGTS TTSSRNARRR ESEKSLENET<br>460 470 480 490 500 510<br>LNKEED**A**HSP TSKPPKPDQP LKVMPAPPPK ENAWVKRSSN PPARSQSSDT EQQSPTSGGG<br>520 530 540 550 560 570<br>KVAPAQPSEE GP**C**RKDENKV DGMNAPKGQT GNSSRGPGDG GNRDHWKESD RKDGKKDQDS<br>580 590 600 610<br>RSAPEPKKPE ENPASKFSSA SKYAALSVDG EDENEGED**C**A E |

| | | |
|---|---|---|
| **DRYG-CTR$_{P332C-W407C}$** | **332-407** | 220 230 240 250 260 270<br>GPRRGDDSF GDKYRDRYDS DRYRDGYRDG YRDGPRRDMD RYGGRDRYDD RGSRDYDRGY<br>280 290 300 310 320 330<br>DSRIGSGRRA FGSGYRRDDD YRGGGDRYED RYDRRDDRSW SSRDDYSRDD YRRDDRGPPQ<br>340 350 360 370 380 390<br>RCKLNLKPRS TPKEDDSSAS TSQSTRAASI FGGAKPVDTA AREREVEERL QKEQEKLQRQ<br>400 410 420 430 440 450<br>LDEPKLERRP RERHPSCRSE ETQERERSRT GSESSQTGTS TTSSRNARRR ESEKSLENET<br>460 470 480 490 500 510<br>LNKEED**A**HSP TSKPPKPDQP LKVMPAPPPK ENAWVKRSSN PPARSQSSDT EQQSPTSGGG<br>520 530 540 550 560 570<br>KVAPAQPSEE GPGRKDENKV DGMNAPKGQT GNSSRGPGDG GNRDHWKESD RKDGKKDQDS<br>580 590 600 610<br>RSAPEPKKPE ENPASKFSSA SKYAALSVDG EDENEGEDYA E |
| **DRYG-CTR$_{W407C-C457}$** | **407-457** | 220 230 240 250 260 270<br>GRRGDDSF GDKYRDRYDS DRYRDGYRDG YRDGPRRDMD RYGGRDRYDD RGSRDYDRGY<br>280 290 300 310 320 330<br>DSRIGSGRRA FGSGYRRDDD YRGGGDRYED RYDRRDDRSW SSRDDYSRDD YRRDDRGPPQ<br>340 350 360 370 380 390<br>RPKLNLKPRS TPKEDDSSAS TSQSTRAASI FGGAKPVDTA AREREVEERL QKEQEKLQRQ<br>400 410 420 430 440 450<br>LDEPKLERRP RERHPSCRSE ETQERERSRT GSESSQTGTS TTSSRNARRR ESEKSLENET<br>460 470 480 490 500 510<br>LNKEEDCHSP TSKPPKPDQP LKVMPAPPPK ENAWVKRSSN PPARSQSSDT EQQSPTSGGG<br>520 530 540 550 560 570<br>KVAPAQPSEE GPGRKDENKV DGMNAPKGQT GNSSRGPGDG GNRDHWKESD RKDGKKDQDS<br>580 590 600 610<br>RSAPEPKKPE ENPASKFSSA SKYAALSVDG EDENEGEDYA E |
| **DRYG-CTR$_{F361C-C457}$** | **361-457** | 220 230 240 250 260 270<br>GRRGDDSF GDKYRDRYDS DRYRDGYRDG YRDGPRRDMD RYGGRDRYDD RGSRDYDRGY<br>280 290 300 310 320 330<br>DSRIGSGRRA FGSGYRRDDD YRGGGDRYED RYDRRDDRSW SSRDDYSRDD YRRDDRGPPQ<br>340 350 360 370 380 390 |

| | | |
|---|---|---|
| | | RPKLNLKPRS TPKEDDSSAS TSQSTRAASI CGGAKPVDTA AREREVEERL QKEQEKLQRQ<br>400 410 420 430 440 450<br>LDEPKLERRP RERHPSWRSE ETQERERSRT GSESSQTGTS TTSSRNARRR ESEKSLENET<br>460 470 480 490 500 510<br>LNKEEDCHSP TSKPPKPDQP LKVMPAPPPK ENAWVKRSSN PPARSQSSDT EQQSPTSGGG<br>520 530 540 550 560 570<br>KVAPAQPSEE GPGRKDENKV DGMNAPKGQT GNSSRGPGDG GNRDHWKESD RKDGKKDQDS<br>580 590 600 610<br>RSAPEPKKPE ENPASKFSSA SKYAALSVDG EDENEGEDYA E |
| **Truncated DRYG-CTR constructs** | | |
| Construct | Labeling sites | Sequence |
| **DRYG** | **-** | 220 230 240 250<br>MKSSHHHHHHENLYFQG RRGDDSF GDKYRDRYDS DRYRDGYRDG YRDGPRRDMD<br>260 270 280 290<br>RYGGRDRYDD RGSRDYDRGY DSRIGSGRRA FGSGYRRDDD<br>300 310 320 330<br>YRGGGDRYED RYDRRDDRSW SSRDDYSRDD YRRDDRGPPQ RC |
| **DRYG-CTR-N** | **332/457** | 220 230 240 250 260 270<br>GRRGDDSF GDKYRDRYDS DRYRDGYRDG YRDGPRRDMD RYGGRDRYDD RGSRDYDRGY<br>280 290 300 310 320 330<br>DSRIGSGRRA FGSGYRRDDD YRGGGDRYED RYDRRDDRSW SSRDDYSRDD YRRDDRGPPQ<br>340 350 360 370 380 390<br>RCKLNLKPRS TPKEDDSSAS TSQSTRAASI FGGAKPVDTA AREREVEERL QKEQEKLQRQ<br>400 410 420 430 440 450<br>LDEPKLERRP RERHPSWRSE ETQERERSRT GSESSQTGTS TTSSRNARRR ESEKSLENET<br>460 470 480 490 500 510<br>LNKEEDC |
| **DRYG-CTR-C** | **609** | 220 230 240 250 260 270 |

| | | |
|---|---|---|
|  |  | GRRGDDSF GDKYRDRYDS DRYRDGYRDG YRDGPRRDMD RYGGRDRYDD RGSRDYDRGY<br>280 290 300 310 320 330<br>DSRIGSGRRA FGSGYRRDDD YRGGGDRYED RYDRRDDRSW SSRDDYSRDD YRRDDRGPPQ<br>R---<br>460 470 480 490 500 510<br>-------HSP TSKPPKPDQP LKVMPAPPPK ENAWVKRSSN PPARSQSSDT EQQSPTSGGG<br>520 530 540 550 560 570<br>KVAPAQPSEE GPGRKDENKV DGMNAPKGQT GNSSRGPGDG GNRDHWKESD RKDGKKDQDS<br>580 590 600 610<br>RSAPEPKKPE ENPASKFSSA SKYAALSVDG EDENEGEDCA E |
| **CTR constructs** | | |
| Construct | Labeling sites | Sequence |
| **CTR$_{P332C\text{-}C457}$** [2] | **332-457** | 340 350 360 370 380 390<br>GCKLNLKPRS TPKEDDSSAS TSQSTRAASI FGGAKPVDTA AREREVEERL QKEQEKLQRQ<br>400 410 420 430 440 450<br>LDEPKLERRP RERHPSWRSE ETQERERSRT GSESSQTGTS TTSSRNARRR ESEKSLENET<br>460 470 480 490 500 510<br>LNKEEDCHSP TSKPPKPDQP LKVMPAPPPK ENAWVKRSSN PPARSQSSDT EQQSPTSGGG<br>520 530 540 550 560 570<br>KVAPAQPSEE GPGRKDENKV DGMNAPKGQT GNSSRGPGDG GNRDHWKESD RKDGKKDQDS<br>580 590 600 610<br>RSAPEPKKPE ENPASKFSSA SKYAALSVDG EDENEGEDYA E |
| **CTR$_{P332C\text{-}W407C}$** | **332-407** | 340 350 360 370 380 390<br>GCKLNLKPRS TPKEDDSSAS TSQSTRAASI FGGAKPVDTA AREREVEERL QKEQEKLQRQ<br>400 410 420 430 440 450<br>LDEPKLERRP RERHPSCRSE ETQERERSRT GSESSQTGTS TTSSRNARRR ESEKSLENET<br>460 470 480 490 500 510<br>LNKEED**A**HSP TSKPPKPDQP LKVMPAPPPK ENAWVKRSSN PPARSQSSDT EQQSPTSGGG |

[2] This CTR construct is used for NMR and CD experiments

| | | |
|---|---|---|
| | | 520 530 540 550 560 570<br>KVAPAQPSEE GPGRKDENKV DGMNAPKGQT GNSSRGPGDG GNRDHWKESD RKDGKKDQDS<br>580 590 600 610<br>RSAPEPKKPE ENPASKFSSA SKYAALSVDG EDENEGEDYA E |
| **CTR$_{F361C-C457}$** | **361-457** | 340 350 360 370 380 390<br>GPKLNLKPRS TPKEDDSSAS TSQSTRAASI CGGAKPVDTA AREREVEERL QKEQEKLQRQ<br>400 410 420 430 440 450<br>LDEPKLERRP RERHPSWRSE ETQERERSRT GSESSQTGTS TTSSRNARRR ESEKSLENET<br>460 470 480 490 500 510<br>LNKEEDCHSP TSKPPKPDQP LKVMPAPPPK ENAWVKRSSN PPARSQSSDT EQQSPTSGGG<br>520 530 540 550 560 570<br>KVAPAQPSEE GPGRKDENKV DGMNAPKGQT GNSSRGPGDG GNRDHWKESD RKDGKKDQDS<br>580 590 600 610<br>RSAPEPKKPE ENPASKFSSA SKYAALSVDG EDENEGEDYA E |
| **CTR$_{W407C-C457}$** | **407-457** | 340 350 360 370 380 390<br>GPKLNLKPRS TPKEDDSSAS TSQSTRAASI FGGAKPVDTA AREREVEERL QKEQEKLQRQ<br>400 410 420 430 440 450<br>LDEPKLERRP RERHPSCRSE ETQERERSRT GSESSQTGTS TTSSRNARRR ESEKSLENET<br>460 470 480 490 500 510<br>LNKEEDCHSP TSKPPKPDQP LKVMPAPPPK ENAWVKRSSN PPARSQSSDT EQQSPTSGGG<br>520 530 540 550 560 570<br>KVAPAQPSEE GPGRKDENKV DGMNAPKGQT GNSSRGPGDG GNRDHWKESD RKDGKKDQDS<br>580 590 600 610<br>RSAPEPKKPE ENPASKFSSA SKYAALSVDG EDENEGEDYA E |
| **CTR$_{C457-G523C}$** | **457-523** | 340 350 360 370 380 390<br>GPKLNLKPRS TPKEDDSSAS TSQSTRAASI FGGAKPVDTA AREREVEERL QKEQEKLQRQ<br>400 410 420 430 440 450<br>LDEPKLERRP RERHPSWRSE ETQERERSRT GSESSQTGTS TTSSRNARRR ESEKSLENET<br>460 470 480 490 500 510<br>LNKEEDCHSP TSKPPKPDQP LKVMPAPPPK ENAWVKRSSN PPARSQSSDT EQQSPTSGGG<br>520 530 540 550 560 570<br>KVAPAQPSEE GPCRKDENKV DGMNAPKGQT GNSSRGPGDG GNRDHWKESD RKDGKKDQDS<br>580 590 600 610 |

| | | |
|---|---|---|
| | | RSAPEPKKPE ENPASKFSSA SKYAALSVDG EDENEGEDYA E |
| **CTR$_{G523C-Y609C}$** | **523-609** | 340 350 360 370 380 390<br>GPKLNLKPRS TPKEDDSSAS TSQSTRAASI FGGAKPVDTA AREREVEERL QKEQEKLQRQ<br>400 410 420 430 440 450<br>LDEPKLERRP RERHPSWRSE ETQERERSRT GSESSQTGTS TTSSRNARRR ESEKSLENET<br>460 470 480 490 500 510<br>LNKEED**A**HSP TSKPPKPDQP LKVMPAPPPK ENAWVKRSSN PPARSQSSDT EQQSPTSGGG<br>520 530 540 550 560 570<br>KVAPAQPSEE GP**C**RKDENKV DGMNAPKGQT GNSSRGPGDG GNRDHWKESD RKDGKKDQDS<br>580 590 600 610<br>RSAPEPKKPE ENPASKFSSA SKYAALSVDG EDENEGED**C**A E |
| **Truncated CTR constructs** | | |
| Construct | Labeling sites | Sequence |
| **CTR-N** | **-** | 340 350 360 370 380 390<br>G**C**KLNLKPRS TPKEDDSSAS TSQSTRAASI FGGAKPVDTA AREREVEERL QKEQEKLQRQ<br>400 410 420 430 440 450<br>LDEPKLERRP RERHPSWRSE ETQERERSRT GSESSQTGTS TTSSRNARRR ESEKSLENET<br>457<br>LNKEED**C** |
| **CTR-C** | **-** | 460 470 480 490 500 510<br>GHSP TSKPPKPDQP LKVMPAPPPK ENAWVKRSSN PPARSQSSDT EQQSPTSGGG<br>520 530 540 550 560<br>KVAPAQPSEE GPGRKDENKV DGMNAPKGQT GNSSRGPGDG GNRDHWKESD<br>570 580 590 600 610<br>RKDGKKDQDS RSAPEPKKPE ENPASKFSSA SKYAALSVDG EDENEGEDYA E |

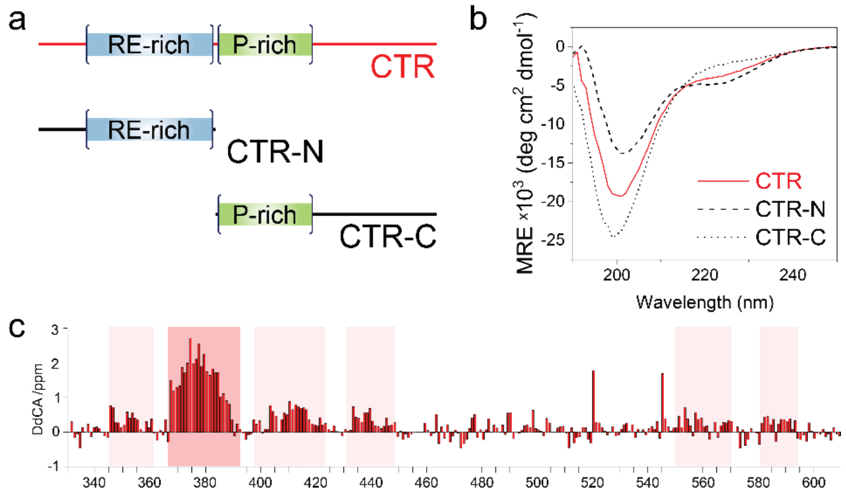

**Supplementary figure 1. Transient helical motifs form in the CTR of eIF4B.** (a) Schematic representation of CTR constructs used. The CTR-N and CTR-C constructs comprise the residues 332-457 and 458-611, respectively. (b) Far-UV CD spectra for full CTR and its N-terminal (CTR-N) and C-terminal (CTR-C) halves. The enhancement of the 220 nm ellipticity signal for CTR-N originates from the formation of a short helical region. (c) Secondary chemical shift values for $^{13}C_{\alpha}$ in CTR, based on random coil values calculated by the online ncIDP server[1]. The regions of increased secondary structure propensity are highlighted in pink. Main helical region is predicted between residues Thr369 and Gln390.

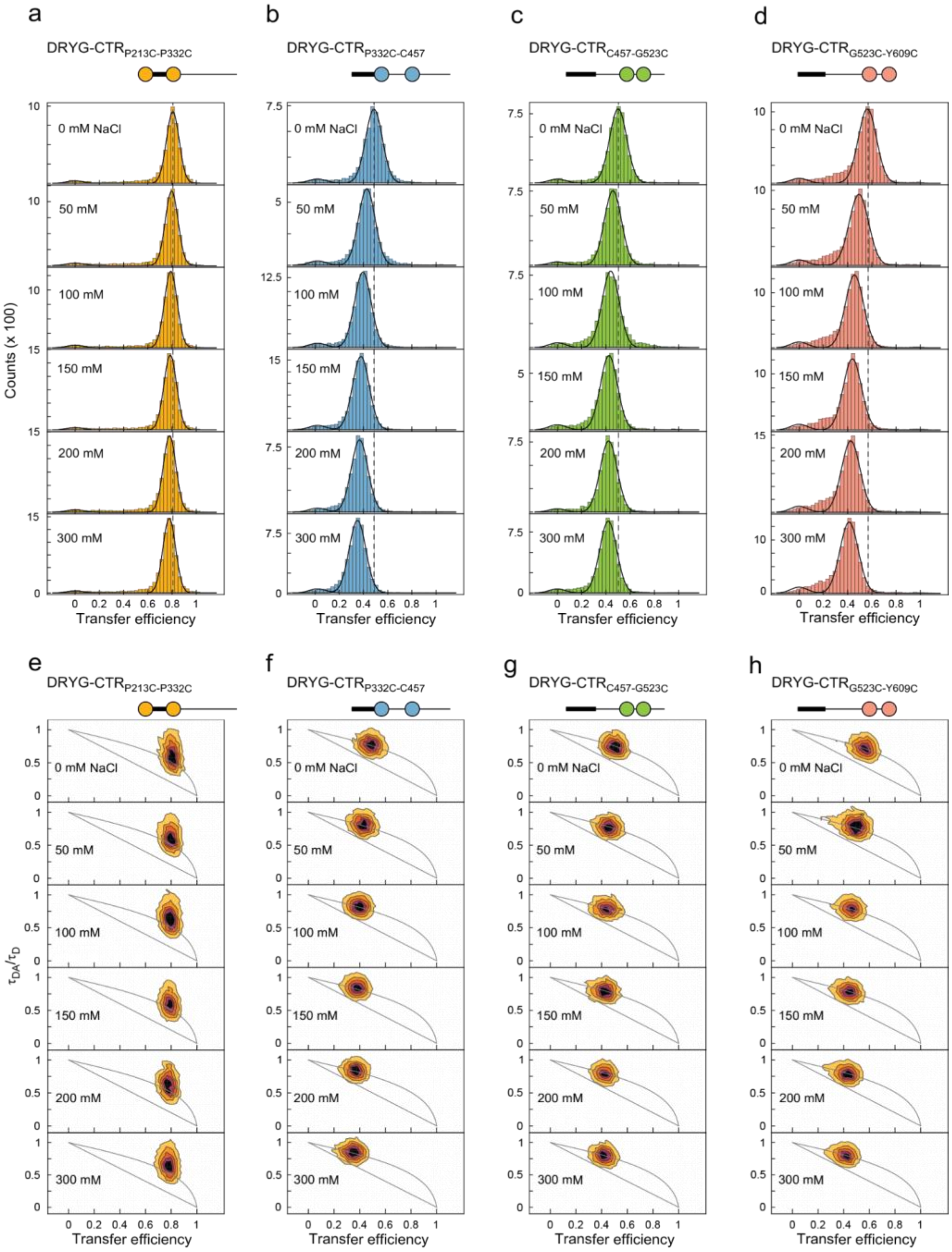

**Supplementary figure 2. Single-molecule FRET analysis of DRYG-CTR constructs at different ionic strengths.** (a-d) Transfer efficiency histograms of DRYG-CTR$_{P213C-P332C}$, DRYG-CTR$_{P332C-C457}$, DRYG-CTR$_{C457-G523C}$ and DRYG-CTR$_{G523C-Y609C}$ in the presence of different concentrations of NaCl. The dashed lines indicate mean transfer efficiency at 0 mM NaCl for respective constructs. (e-h) Normalized fluorescence lifetime - transfer efficiency 2D histograms for each construct under the same conditions as transfer efficiency histograms.

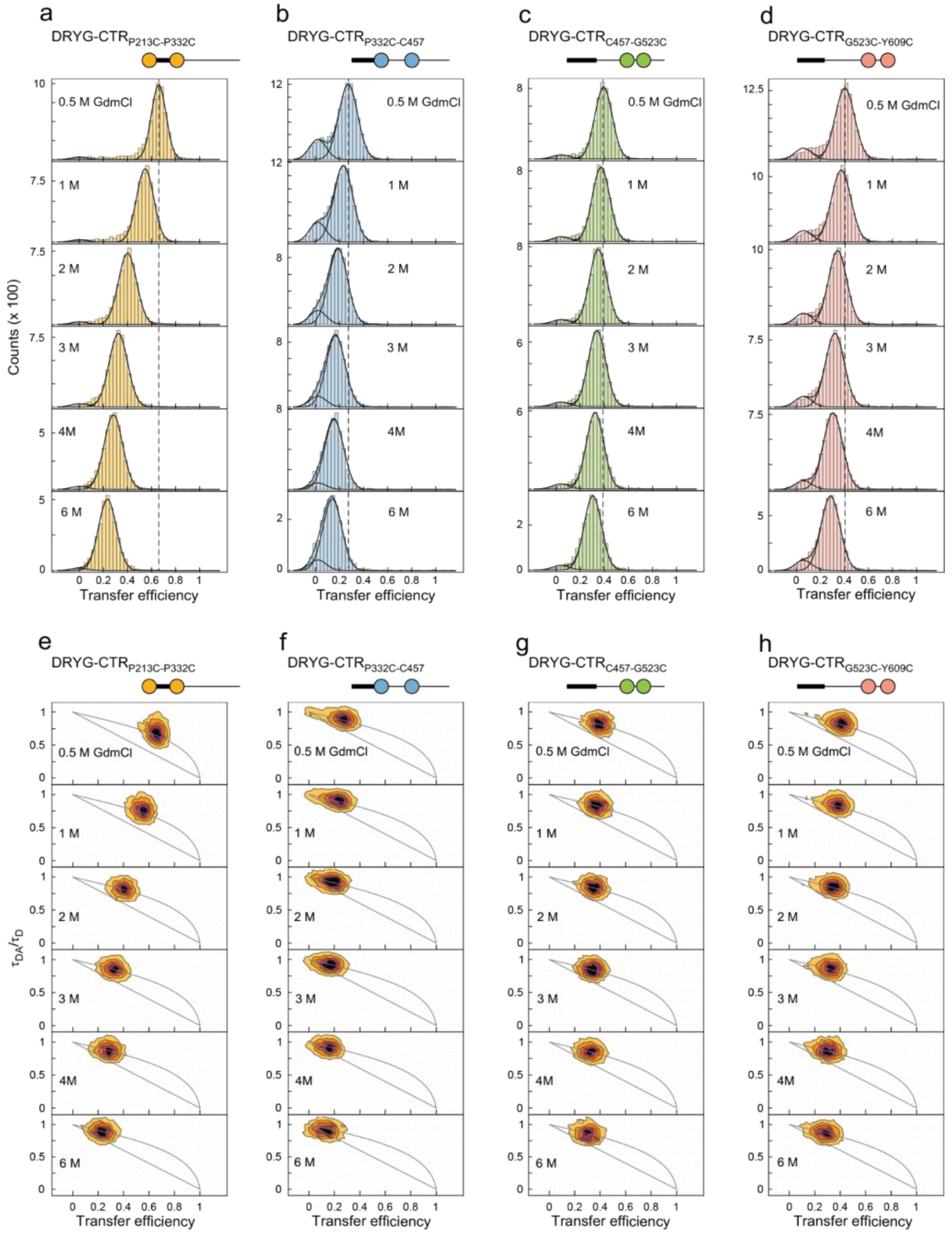

**Supplementary figure 3. Single-molecule FRET analysis of DRYG-CTR constructs in the presence of GdmCl.** (a-d) Transfer efficiency histograms of DRYG-CTR$_{P213C\text{-}P332C}$, DRYG-CTR$_{P332C\text{-}C457}$, DRYG-CTR$_{C457\text{-}G523C}$ and DRYG-CTR$_{G523C\text{-}Y609C}$ in the presence of different concentrations of GdmCl. The dashed lines indicate mean transfer efficiency at 0.5 M GdmCl for respective constructs. (e-h) Normalized fluorescence lifetime - transfer efficiency 2D histograms for each construct under the same conditions as transfer efficiency histograms.

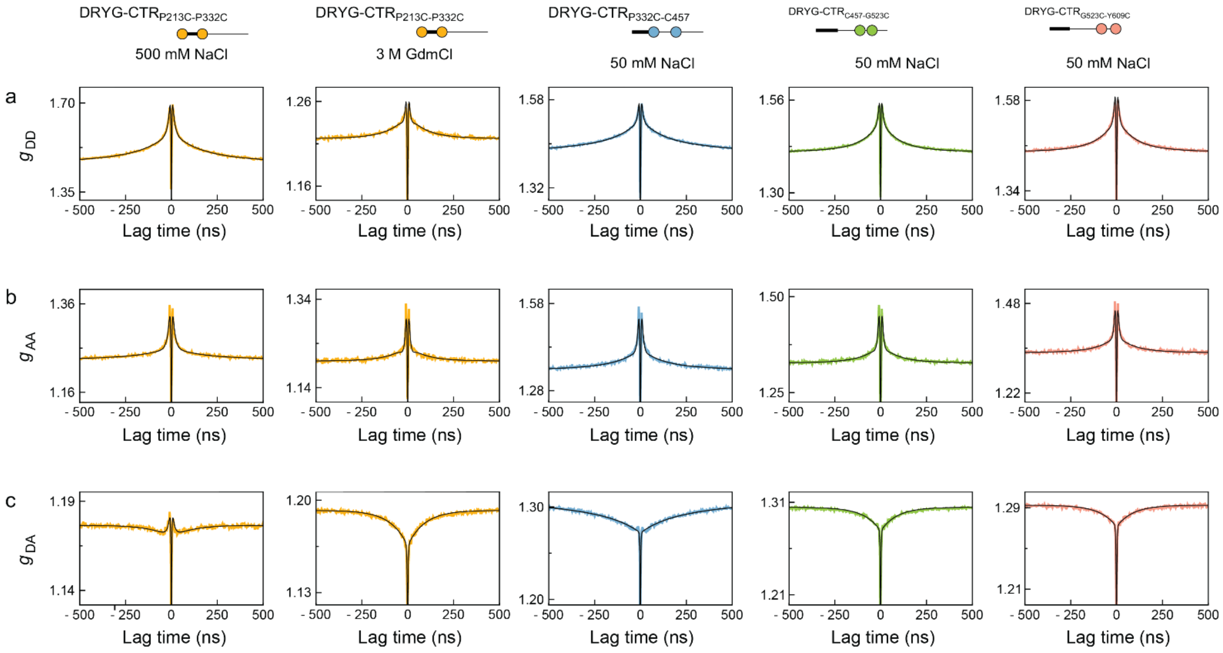

**Supplementary figure 4. nsFCS analysis of DRYG-CTR constructs at different conditions as in Fig. 3d.** Donor-donor (a), acceptor-acceptor (b) and donor-acceptor (c) correlation curves for DRYG-CTR$_{P213C-P332C}$ acquired at 500 mM NaCl and 3 M GdmCl, and for the DRYG-CTR$_{P332C-C457}$, DRYG-CTR$_{C457-G523C}$ and DRYG-CTR$_{G523C-Y609C}$ at 50 mM NaCl. The solid lines are global fits using Eq. 7.

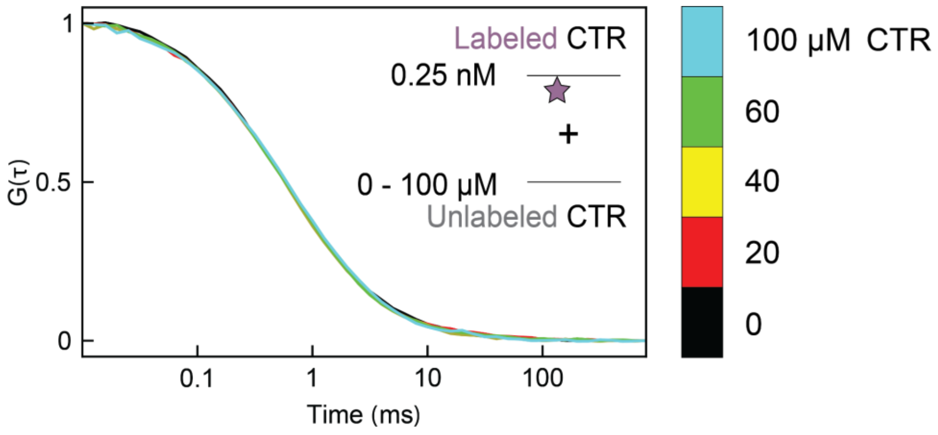

**Supplementary figure 5. FCS data for CTR self-association.** A set of FCS curves obtained at increasing concentrations of unlabeled protein show no change in autocorrelation curve of CTR, demonstrating an absence of CTR-CTR self-association at 50 mM NaCl. The insert shows a schematic of FCS oligomerization assay, based on a titration of sub-nanomolar labeled CTR with micromolar concentrations of unlabeled CTR protein.

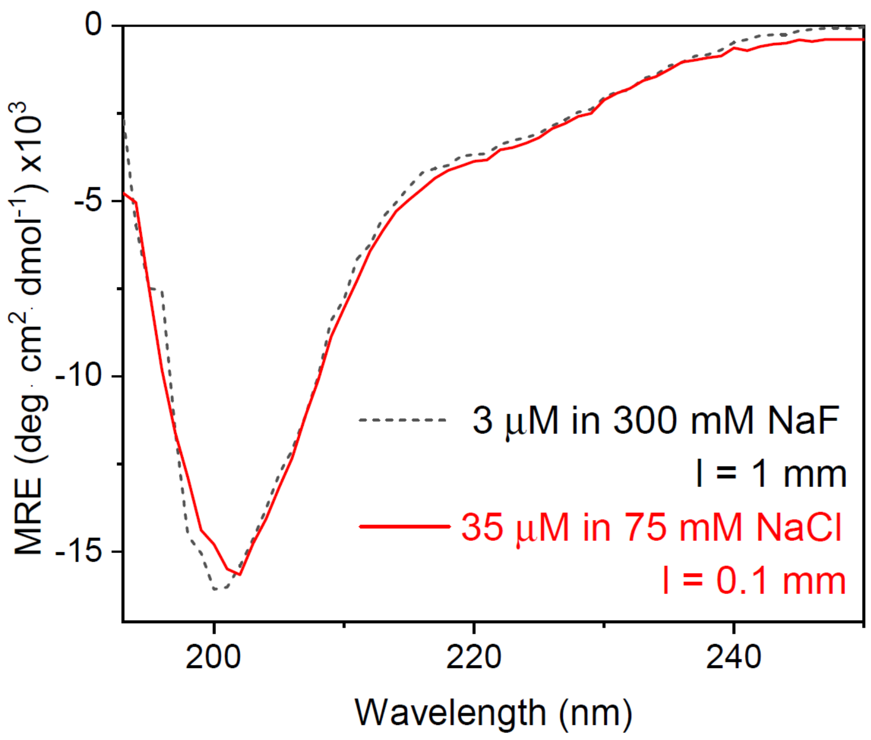

**Supplementary figure 6. CD spectra reveal no change in secondary structure content upon eIF4B IDR oligomerization.** The spectrum of 3 µM DRYG-CTR solutions (dashed black) in 20 mM NaP, 300 mM NaF, 1mM TCEP pH 7.0 buffer measured in 1 mm pathlength cuvette and the spectrum of 35 µM DRYG-CTR solutions (solid red) in 20 mM NaP, 75 mM NaCl, 1mM TCEP pH 7.0 buffer measured in 0.1 mm pathlength cuvette.

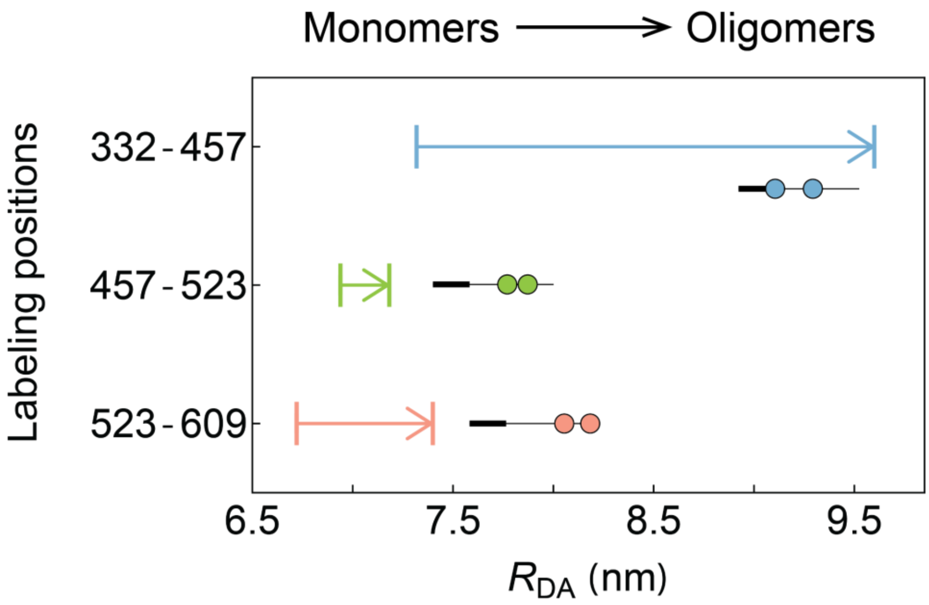

**Supplementary figure 7. The eIF4B CTR undergoes expansion upon oligomerization.** The change of inter-residue distance for DRYG-CTR$_{P332C-C457}$, DRYG-CTR$_{C457-G523C}$ and DRYG-CTR$_{G523C-Y609C}$ constructs, showing an expansion of corresponding protein regions upon protein self-association.

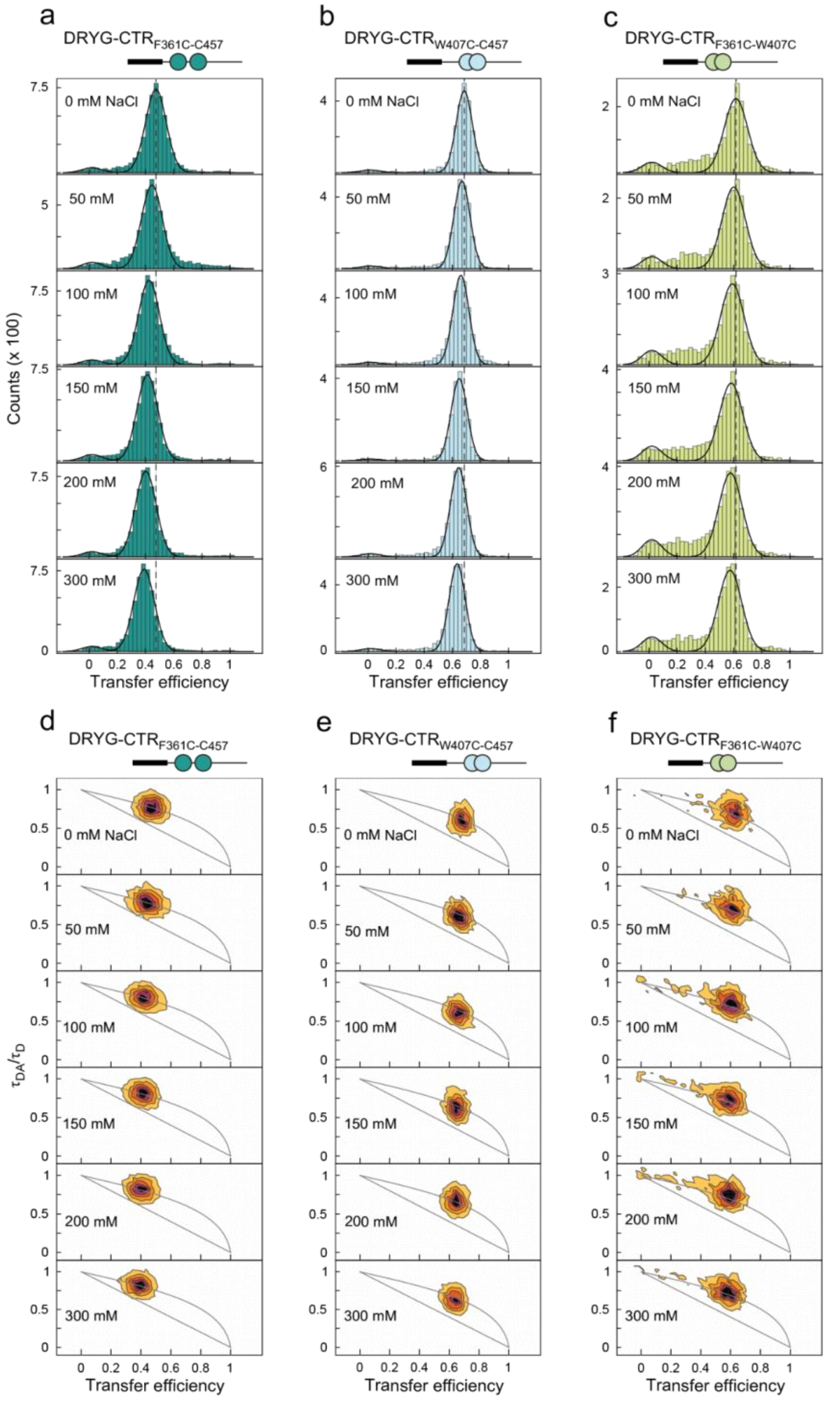

**Supplementary figure 8. Single-molecule FRET analysis of additional DRYG-CTR constructs at different ionic strengths.** (a-c) Transfer efficiency histograms of DRYG-CTR$_{F361C-C457}$, DRYG-CTR$_{W407C-C457}$ and DRYG-CTR$_{F361C-W407C}$ in the presence of different concentrations of NaCl. The dashed lines indicate mean transfer efficiency at 0 mM NaCl for respective constructs. (d-f) Normalized fluorescence lifetime - transfer efficiency 2D histograms for each construct under the same conditions as transfer efficiency histograms.

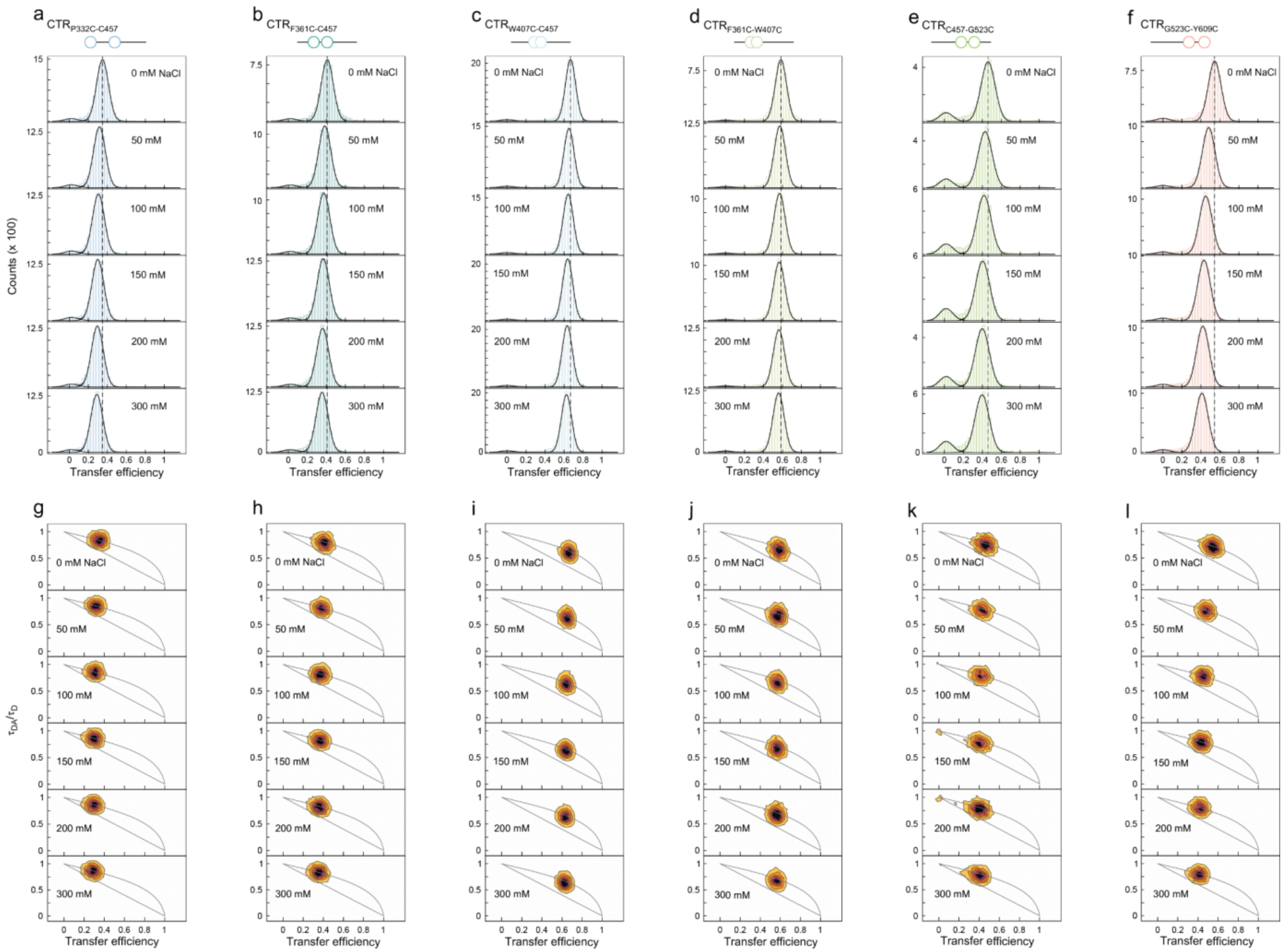

**Supplementary figure 9. Single-molecule FRET analysis of CTR constructs at different ionic strengths.** (a-f) Transfer efficiency histograms of $CTR_{P332C\text{-}C457}$, $CTR_{F361C\text{-}C457}$, $CTR_{W407C\text{-}C457}$, $CTR_{F361C\text{-}W407C}$, $CTR_{C457\text{-}G523C}$ and $CTR_{G523C\text{-}Y609C}$ in the presence of different concentrations of NaCl. The dashed lines indicate mean transfer efficiency at 0 mM NaCl for respective constructs in respective figures. (g-l) Normalized fluorescence lifetime - transfer efficiency 2D histograms for each construct under the same conditions as transfer efficiency histograms.

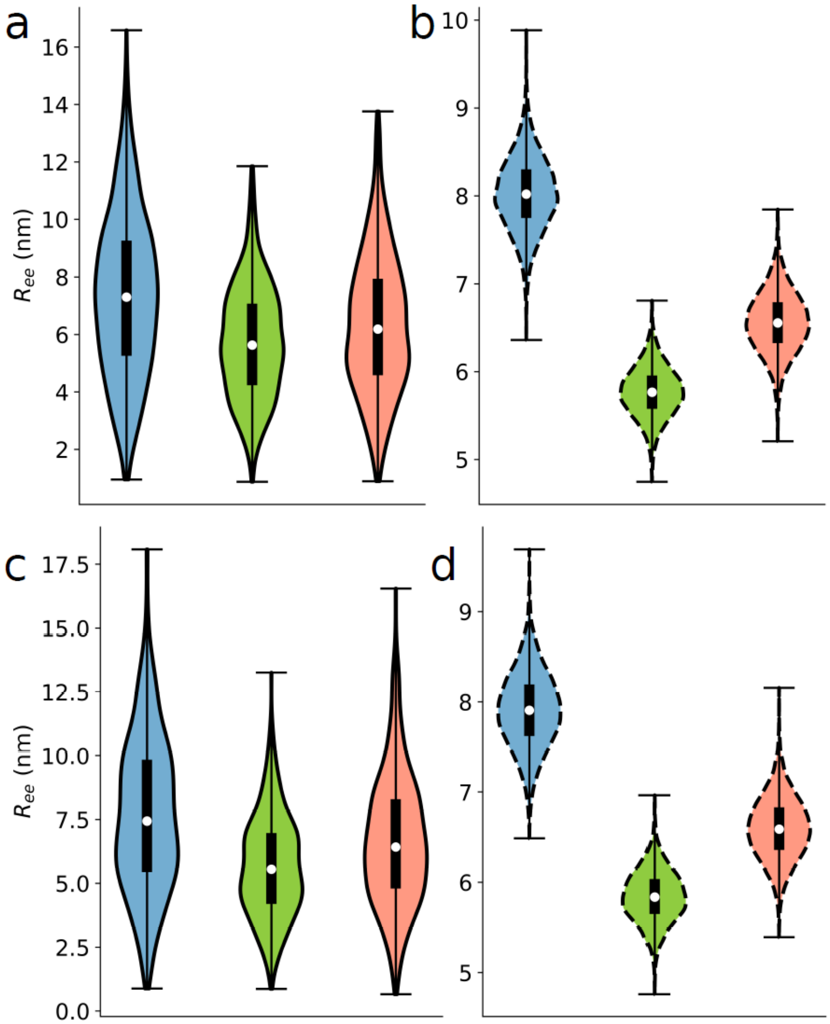

**Supplementary figure 10: End-to-end distance distributions of the experimentally investigated eIF4B constructs.** (a-b) End-to-end distances ($R_{ee}$) between residues 332-457 (blue), 457-523 (green), and 523-609 (pink), within monomeric (solid black line) and oligomeric DRYG-CTR (dashed black line), at an ionic strength of $I$ = 117 mM, as in experiments. (c-d) End-to-end distances ($R_{ee}$) between residues 332-457 (blue), 457-523 (green), and 523-609 (pink), for monomeric DRYG-CTR (solid black line) and oligomeric DRYG-CTR (dashed black line), simulated at an ionic strength of $I$ = 192 mM.

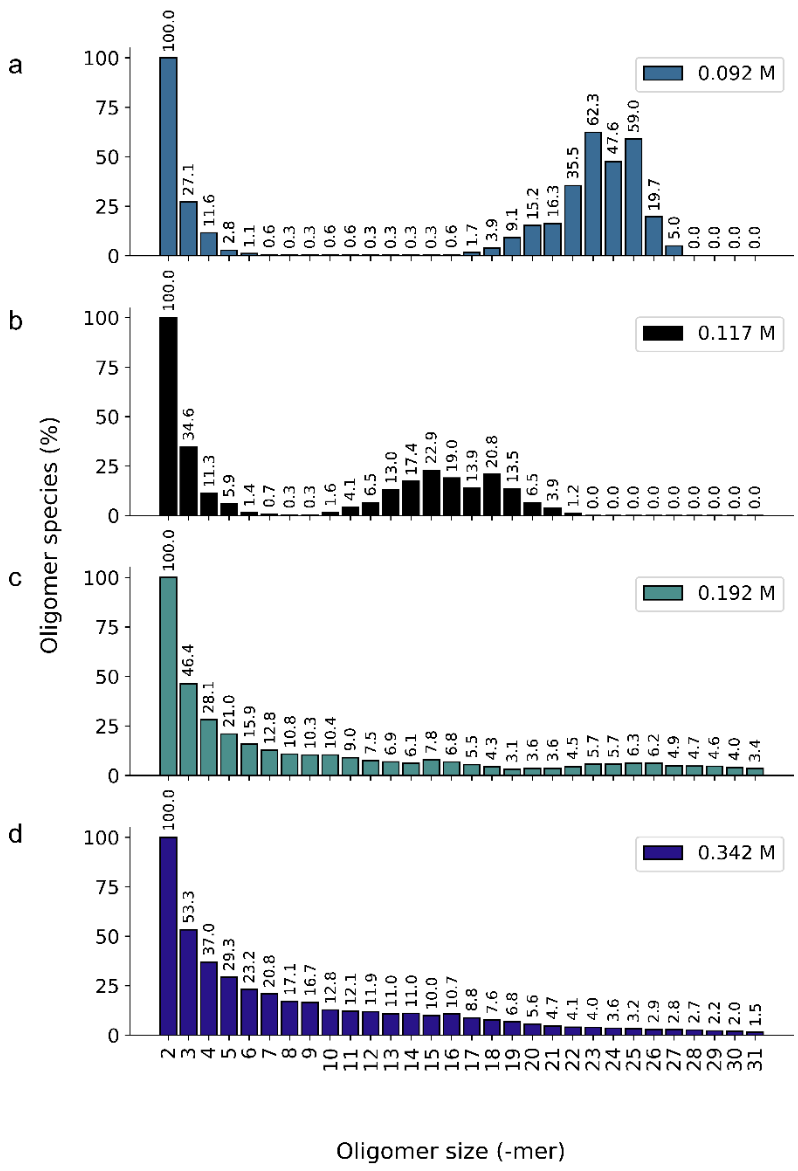

**Supplementary figure 11: The size distribution and relative abundances of DRYG-CTR oligomers at different ionic strengths.** Bar plot showing the abundance of the 30 most sampled oligomeric species obtained from the simulation of DRYG-CTR *I* = 92, 117, 192, 342 mM (a-d) ionic strength. Each oligomeric size is expressed as a percentage of the simulated trajectory and normalized based on the size of the smallest oligomer (2-mer).

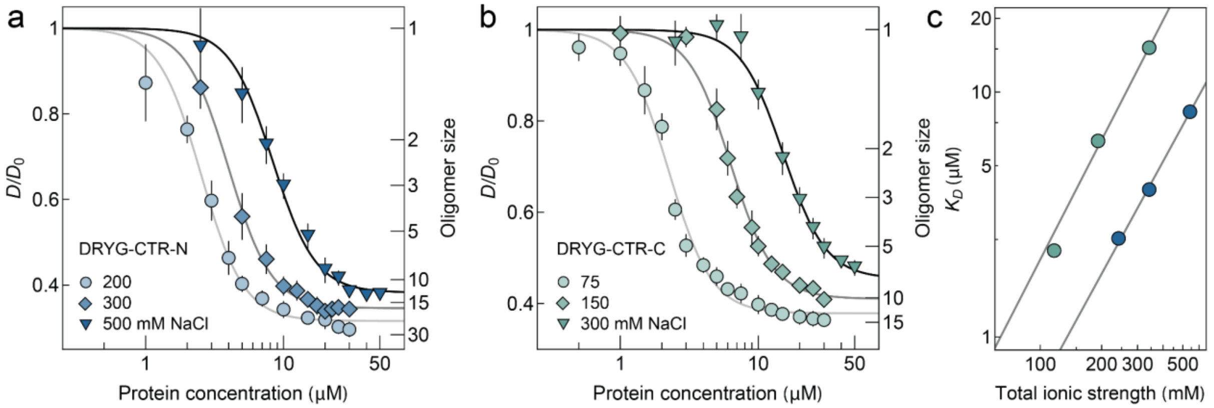

**Supplementary figure 12. The self-association of truncated DRYG-CTR constructs.** The dependence of relative diffusion rates of DRYG-CTR-N (a) and DRYG-CTR-C (b) on protein concentration, acquired in buffers containing various concentrations of NaCl, indicates a systematic reduction of protein association affinity with increasing buffer ionic strength. The solid lines show global fits to the Hill equation. (d) The dependence of apparent dissociation constants of DRYG-CTR-N and DRYG-CTR-C oligomers on buffer ionic strength. The black solid line is a fit to $\log K_D = \log K_D^{1M} - \Delta n \cdot \log c$ equation.

```
1          11         21         31         41
MAASAKKKNK KGKTISLTDF LAEDGGTGGG STYVSKPVSW ADETDDLEGD
eeeeeeeeee eeeebebbeb bbeeeeeeee eeeeeeeeee eeeeeebeee
ff    f  f f ff fs fs ss f            f f   ffffffs f

51         61         71         81         91
VSTTWHSNDD DVYRAPPIDR SILPTAPRAA REPNIDRSRL PKSPPYTAFL
ebbeeeeeee ebeebeebee ebbeebeeeb eeeeeeeeeb eeeeebbbbb
  s  ff       f    f  f sffsf fs  ff      s f   f ssss

101        111        121        131        141
GNLPYDVTEE SIKEFFRGLN ISAVRLPREP SNPERLKGFG YAEFEDLDSL
eebeeebeee bbeebbebbb bbbbebeeee eeeeebeebb ebebeebeeb
ff ff s f  ss          s s    f   ff   f      f ff  f

151        161        171        181        191
LSALSLNEES LGNRRIRVDV ADQAQDKDRD DRSFGRDRNR DSDKTDTDWR
bebbeeeeee eeeeebebbb beeeeeeeee eeeeeeeeee eeeeeeeebe
     f ff    fffs ss  sfff  f f      f ff     f ff f f

201        211        221        231        241
ARPATDSFDD YPPRRGDDSF GDKYRDRYDS DRYRDGYRDG YRDGPRRDMD
beeeeeebee eeeeeeeeee eeebeeeeee eeeeeeeeee eeeeeeeeee
s            f f               f f

251        261        271        281        291
RYGGRDRYDD RGSRDYDRGY DSRIGSGRRA FGSGYRRDDD YRGGGDRYED
eeeeeeeeee eeeeeeeeee ebebeebeeb ebebeeeeee eeeeeeeeee
         f       ff    s     ff  f     fff   f

301        311        321        331        341
RYDRRDDRSW SSRDDYSRDD YRRDDRGPPQ RPKLNLKPRS TPKEDDSSAS
eeeeeeeeee eeeeeeeeee eeeeeeeeee eeebeeeeee eeeeeeeeee
                           f   f ff  f  f f

351        361        371        381        391
TSQSTRAASI FGGAKPVDTA AREREVEERL QKEQEKLQRQ LDEPKLERRP
eeeeeebbbb beebeeeeee beeeebeeeb eeeeeebeee eeeeeeeeee
        s      f ffff  ffffsff s ff       f        f

401        411        421        431        441
RERHPSWRSE ETQERERSRT GSESSQTGTS TTSSRNARRR ESEKSLENET
eeeeeebeee eeeeeeeeee eeeeeeeeee eeeeeeeeee eeeeeeeeee
 f   f             ff  ffff                     f  f

451        461        471        481        491
LNKEEDCHSP TSKPPKPDQP LKVMPAPPPK ENAWVKRSSN PPARSQSSDT
beeeeeeeee eeeeeeeeee beebeeeeee eebebeeeee eeeeeeeeee
        f              ff ffffff ff   f

501        511        521        531        541
EQQSPTSGGG KVAPAQPSEE GPGRKDENKV DGMNAPKGQT GNSSRGPGDG
eeeeeeeeee eeeeeeeeee eeeeeeeeee eeeeeeeeee eeeeeeeeee
                            f

551        561        571        581        591
GNRDHWKESD RKDGKKDQDS RSAPEPKKPE ENPASKFSSA SKYAALSVDG
eeeeeeeeee eeeeeeeeee eeeeeeeeee eeeeeebeeb bebbbbbbee
                                 f      f    f  s   f

601        611
EDENEGEDYA E
eeeeeeeeee e
```

e - An exposed residue according to the neural-network algorithm.
b - A buried residue according to the neural-network algorithm.
f - A predicted functional residue (highly conserved and exposed).
s - A predicted structural residue (highly conserved and buried).

**Supplementary figure 13. The prediction of eIF4B sequence conservation according to ConSurf[2].** The conservation scale is 1 to 9, with 1-4 for variable, 5 for average and 6-9 conserved according to the following color code: 1 2 3 4 5 6 7 8 9. The symbols below refer to the categories shown in the box on the right side of the sequence.

References

1 Tamiola, K., Acar, B. & Mulder, F. A. Sequence-specific random coil chemical shifts of intrinsically disordered proteins. *J Am Chem Soc* **132**, 18000-18003 (2010). https://doi.org:10.1021/ja105656t

2 Ashkenazy, H. *et al.* ConSurf 2016: an improved methodology to estimate and visualize evolutionary conservation in macromolecules. *Nucleic Acids Res* **44**, W344-350 (2016). https://doi.org:10.1093/nar/gkw408
